# The NAD^+^ precursor NMN activates dSarm to trigger axon degeneration in *Drosophila*

**DOI:** 10.1101/2022.01.30.478002

**Authors:** Arnau Llobet Rosell, Maria Paglione, Jonathan Gilley, Magdalena Kocia, Massimiliano Gasparrini, Lucia Cialabrini, Nadia Raffaelli, Carlo Angeletti, Giuseppe Orsomando, Michael P. Coleman, Andrea Loreto, Lukas J. Neukomm

## Abstract

Axon degeneration contributes to the disruption of neuronal circuit function in diseased and injured nervous systems. Severed axons degenerate following the activation of an evolutionarily conserved signaling pathway, which culminates in the activation of SARM1 in mammals to execute the pathological depletion of the metabolite NAD^+^. SARM1 NADase activity is activated by the NAD^+^ precursor nicotinamide mononucleotide (NMN). In mammals, keeping NMN levels low potently preserves axons after injury, however, it remains unclear whether NMN is also a key mediator of axon degeneration, and dSarm activation, in flies. Here, we demonstrate that lowering NMN levels in *Drosophila* through the expression of a newly generated prokaryotic NMN-Deamidase (NMN-D) preserves severed axons for months and keeps them circuit-integrated for weeks. NMN-D alters the NAD^+^ metabolic flux by lowering NMN, while NAD^+^ remains unchanged *in vivo*. Increased NMN synthesis, by the expression of mouse nicotinamide phosphoribosyltransferase (mNAMPT), leads to faster axon degeneration after injury. We also show that NMN-induced activation of dSarm mediates axon degeneration *in vivo*. Finally, NMN-D delays neurodegeneration caused by loss of the sole NMN-consuming and NAD^+^-synthesizing enzyme dNmnat. Our results reveal a critical role for NMN in neurodegeneration in the fly, which extends beyond axonal injury. The potent neuroprotection by reducing NMN levels is similar or even stronger than the interference with other essential mediators of axon degeneration *in Drosophila*.

## Introduction

The elimination of large portions of axons is a widespread event in the developing nervous system ^1,2^. Axon degeneration is also an early hallmark of nervous system injury and a common feature of neurodegenerative diseases ^3–5^. Understanding the underlying molecular mechanisms may facilitate the development of treatments to block axon loss in acute or chronic neurological conditions.

Wallerian degeneration is a well-established, evolutionarily conserved, and simple system to study how injured axons execute their self-destruction ^6,7^. Upon axonal injury (axotomy), distal axons separated from their soma degenerate within a day. Axotomy activates a signaling pathway (programmed axon degeneration, or axon death) that actively executes the self-destruction of severed axons. Induced signaling culminates in the activation of dSarm in flies and SARM1 in mice ^8,9^. As an NADase, once activated, dSarm/SARM1 executes the pathological depletion of nicotinamide adenine dinucleotide (NAD^+^) in severed axons, culminating in catastrophic fragmentation ^10–12^. Initially thought to be activated only after injury, over recent years it became evident that axon death signaling is also activated in many non-injury neurological disorders^13,14^.

In mammals, SARM1 activation is tightly controlled by metabolites in the NAD^+^ biosynthetic pathway. Axonal transport ensures continuous delivery of the labile enzyme nicotinamide mononucleotide adenylyltransferase 2, NMNAT2^15^, an axonal survival factor that consumes nicotinamide mononucleotide (NMN) to synthesize NAD^+^. When axons are injured, NMNAT2 is rapidly degraded by the E3 ubiquitin ligase PHR1 and MAPK signaling ^16,17^. This leads to a temporal rise of axonal NMN, whereas NAD^+^ biosynthesis halts ^18–21^. NMN and NAD^+^ compete by binding to an allosteric pocket in the SARM1 armadillo-repeat (ARM) domain, crucial for SARM1 activation. While a rise in NMN induces a conformational change to activate SARM1 ^11,22,23^, NAD^+^ prevents this activation by competing for the same pocket in the ARM domain^24^ and this competitive binding occurs at physiologically relevant levels of both NMN and NAD^+^ ^25^.

Previous studies have shown that expression of the prokaryotic enzyme PncC, also known as NMN-Deamidase (NMNd), which converts NMN to nicotinic acid mononucleotide (NaMN) ^26^, prevents SARM1 activation and preserves severed axons in mammals and zebrafish: for instance, axons with NMNd remain preserved up to 96 h in murine neuronal cultures ^18,19,27^, in zebrafish up to 24 h and mice up to 3 weeks^20^. It remains currently unclear whether NMNd expression levels determine the extent of preservation.

Much of this mechanism of axon death signaling is conserved in *Drosophila* However, flies harbor some noticeable differences. A single *dnmnat* gene provides a cytoplasmic and nuclear splice variant; *dnmnat* interference results therefore in cellular dNmnat loss ^28^. dNmnat turnover is regulated solely by the E3 ubiquitin ligase Hiw ^29^, but not by MAPK signaling ^30^. And the BTB/Back domain-containing Axed executes catastrophic fragmentation downstream of NAD^+^ depletion, while the mammalian paralog(s) remain to be identified ^30^.

The role of NMN in axon degeneration in *Drosophila* is also controversial. Flies lack nicotinamide phosphoribosyltransferase (NAMPT) ^31^. NMN might therefore not be a prominent intermediate in the NAD^+^ biosynthetic pathway, thus having a minor role as mediator of axon degeneration ^32^. The Gal4/UAS-mediated NMNd expression in *Drosophila* neurons preserves severed axons for 3–5 days after injury ^33^. It contrasts the phenotype of other axon death signaling mediators, such as loss-of-function mutations in *hiw*, *dsarm*, and *axed*, as well as over-expression of *dnmnat* (*dnmnat^OE^*), all of which harbor severed axons that remain preserved for weeks to months ^9,30,34,35^. Therefore, the role of NMN as an axon death mediator in *Drosophila* remains to be fully determined.

Here, we report that NMN is a key mediator of injury-induced axon degeneration *in Drosophila*. Genetic modifications resulting in low NMN levels protect severed axons for the lifespan of the fly, while the addition of an extra NMN synthesizing activity forces axons to undergo faster degeneration after injury. NMN induces the dSarm NADase activity, demonstrating its role as a crucial activator *in vivo*.

## Results

### Robust expression of prokaryotic NMN-Deamidase in *Drosophila*

Mutations in *hiw, dsarm*, or *axed* attenuate axon death signaling resulting in morphological preservation of severed axons for approximately 50 days, the average lifespan of *Drosophila* ^9,30,35^. In contrast, neuronal expression of prokaryotic NMN-Deamidase (NMNd) to consume NMN results in less than 10 % of severed axons being preserved at 7 days post axotomy (dpa) ^33^. We performed an established wing injury assay ^36^ to confirm this observation. Briefly, a subset of GFP-labeled sensory neurons (e.g., *dpr1-Gal4* MARCM clones) expressing NMNd ^33^ or GFP were subjected to axotomy in 5–7-day aged flies with one wing being partially injured and the other serving as an uninjured control (Fig. S1A). At 7 dpa, we quantified uninjured control axons, axonal debris and severed intact axons, respectively (Fig. S1B, left), and calculated the percentage of protected severed axons (Fig. S1B, right). We observed a 40 % preservation of severed axons with NMNd (Fig. S1, genotypes in Table S1). This is probably due to higher NMNd levels, because of higher Gal4 levels in *dpri* than *elaV* ^33^. However, given that the expression of NMNd fails to attenuate axon death signaling to the extent of axon death mutants suggests that either, NMN does not play an essential role in activating axon death supported by the absence of the NMN-synthesizing enzyme Nampt in flies ^31^, or NMNd expression and the resulting NMN consumption is simply not sufficient for potent attenuation of axon death signaling and preservation of severed axons.

Based on the above observations, we decided to generate an N-terminal GFP-tagged NMN-Deamidase to increase protein stability ^37^, which we named GFP::NMN-D. Plasmids with GFP-tagged wild-type and enzymatically dead versions of NMN-Deamidase were generated ^19,20^, under the control of the UAS regulatory sequence (*UAS–GFP::NMN-D*, and *UAS–GFP::NMN-D^dead^*, respectively). We found that both wild-type and enzymatically dead NMN-D enzymes are equally expressed in S2 cells, as detected by newly generated PncC antibodies (Fig. S2A). Notably, we observed two immunoreactivities per lane, with the lower band being a potential degradation product.

The equal expression of the NMN-D enzymes prompted us to use the plasmids to generate transgenic flies by targeted insertion (*attP40* landing site). To compare *in vivo* expression levels, NMN-D, NMN-D^dead^, and NMNd were expressed with pan-cellular *actin–Gal4*. We found that NMN-D and NMN-D^dead^ immunoreactivities were significantly stronger than NMNd (Fig. S2B, C). In addition, GFP immunoreactivity was also detected in NMN-D and NMN-D^dead^ confirming the robust expression of the newly generated tagged proteins (Fig.1A). These results show that our newly generated GFP-tagged NMN-D variants are substantially stronger expressed than NMNd *in vivo*.

**Figure 1.**
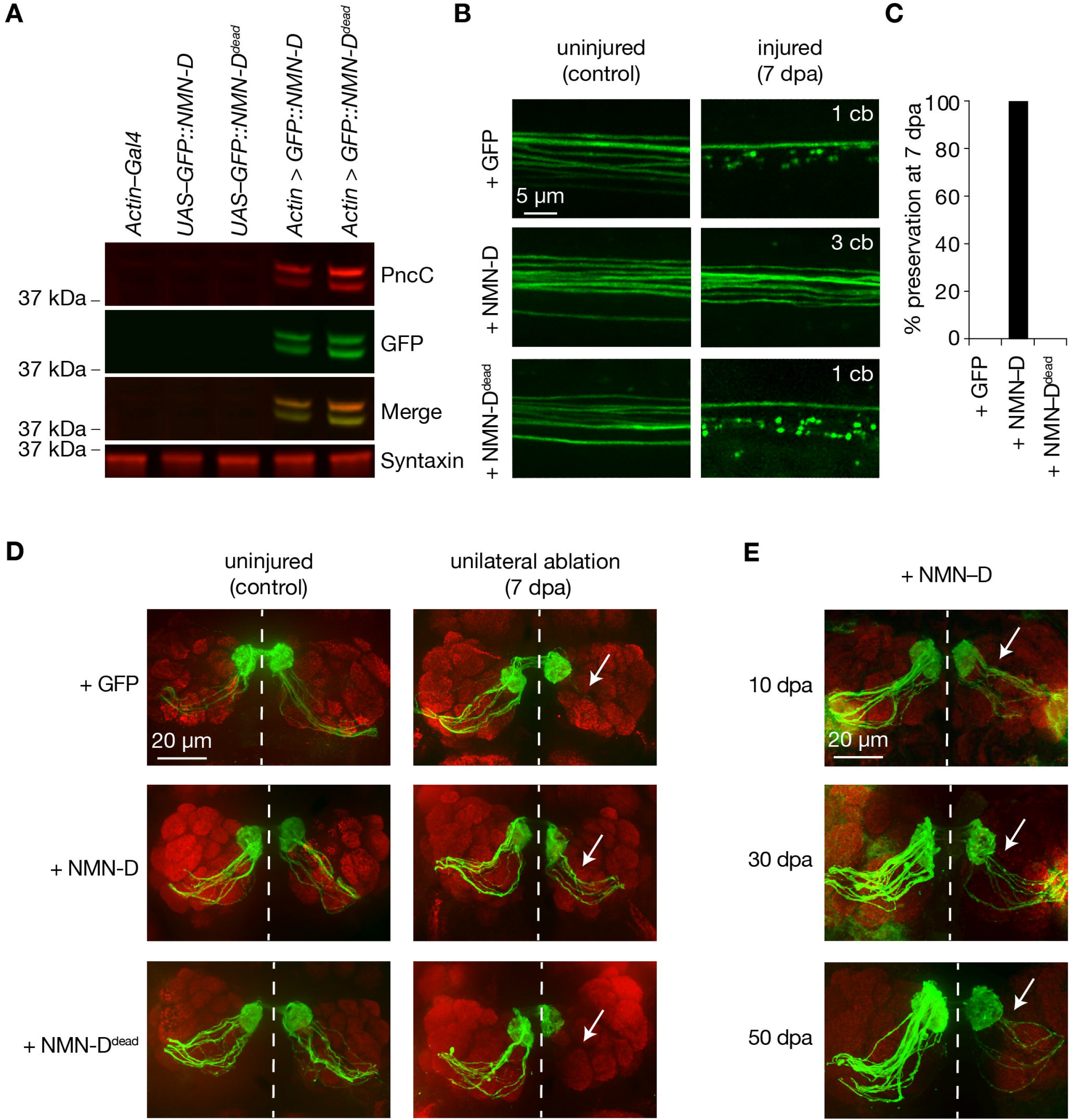
Neuronal expression of prokaryotic NMN-D preserves the morphology of severed axons for the lifespan in *Drosophila*. **A** Equal expression of wild-type and enzymatic dead NMN-D enzymes, respectively. Western blots with anti-PncC and anti-GFP immunoreactivities (red and green, respectively). **B** Low NMN results in severed wing sensory neuron axons that remain morphologically preserved at 7 dpa. Examples of control and 7 dpa. **C** Axon death quantification. % preservation of injured axons at 7 dpa, average ± standard error of the mean (*n* = 15 wings). D Low NMN results in severed olfactory receptor neuron axons that remain morphologically preserved at 7 dpa. Examples of control and 7 dpa (arrows, site of unilateral ablation). E Low NMN results in severed axons that remain morphologically preserved for 50 days. Examples of 10, 30, and 50 dpa (arrows, site of unilateral ablation).

### Neuronal NMN-D expression blocks injury-induced axon degeneration for the lifespan of *Drosophila*

The lower-expressed NMNd resulted in a 40 % preservation in our wing injury assay. We repeated the injury assay to assess the preservation of our newly generated and higher-expressed NMN-D variants. While severed axons with GFP or NMN-D^dead^ degenerated, axons with NMN-D remained fully preserved at 7 dpa (Fig. 1B, C, S3). This contrasts with the weaker preservation of axons with lower NMNd expression (Fig. S1). Similarly, strong preservation was seen in cholinergic olfactory receptor neurons (ORNs), where severed axons with NMN-D remained preserved at 7 dpa (Fig. 1D). We extended the ORN injury assay and found preservation at 10, 30, and 50 dpa (Fig. 1E). Thus, high levels of NMN-D confer robust protection of severed axons for multiple neuron types for the entire lifespan of *Drosophila*.

### NMN-D alters the NAD^+^ metabolic flux to lower NMN in heads

Before measuring the effect of NMN-D on the NAD^+^ metabolic flux, we measured the activities of the various NAD^+^ biosynthetic enzymes in fly heads (Tab. S2, Fig. 2A) ^38,39^. We demonstrated that NAD^+^ can be synthesized from quinolinate, nicotinamide (NAM) and nicotinamide riboside (NR). We also confirmed the absence of NAMPT activity^31^. Next, we wanted to know whether the sole expression of NMN-D can change the NAD^+^ metabolic flux *in vivo* (Fig. 2A). We compared levels of metabolites in heads by LC-MS/MS among samples that expressed NMN-D and NMN-D^dead^ (Fig. 2B) ^40^. Consistent with robust NMN-D activity, NMN levels were 6-times lower and NaMN 12-times higher. We also found significantly higher NaAD and NaR levels. Importantly, all other metabolites remained unchanged, including NAD^+^ (Fig. 2B). These observations demonstrate that the sole expression of our newly generated NMN-D is sufficient to change the NAD^+^ metabolic flux, thereby significantly lowering NMN levels without affecting NAD^+^ in *Drosophila* heads; they serve as an excellent tissue for metabolic analyses.

**Figure 2.**
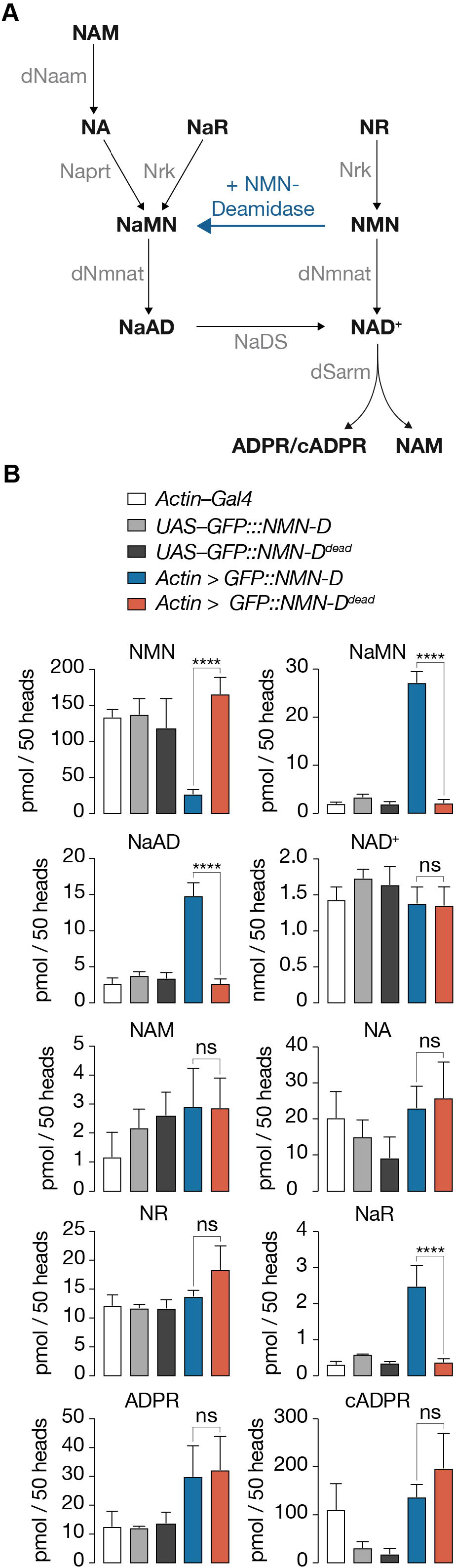
Pan-cellular NMN-D expression alters the flux of NAD^+^ metabolites to lower NMN in heads of *Drosophila*. **A** *Drosophila* NAD^+^ metabolic pathway. Black, metabolites; grey, enzymes; blue, prokaryotic NMN-D, respectively. B The expression of NMN-D results in lower NMN and higher NaMN, NaAD, and NaR levels, respectively. Extracted metabolites from 50 heads, mean ± standard deviation (*n =* 4). One-way ANOVA with Tuckey’s multiple comparisons test, **** = *p* < 0.0001, ns = *p* > 0.05.

### Low axonal NMN preserves synaptic connectivity for weeks after injury

Mutations that attenuate axon death signaling exhibit robust suppression of morphological degeneration after axotomy. They also preserve synaptic connectivity. We have previously demonstrated that synaptic connectivity of severed axons with attenuated axon death remains preserved for at least 14 days, using a simple optogenetic assay ^30,36^. Briefly, mechanosensory chordotonal neurons in the Johnston’s organ (JO), whose cell bodies are in the 2^nd^ segment of adult antennae, are required and sufficient for antennal grooming ^41,42^. The JO-specific expression of CsChrimson combined with a red-light stimulus is sufficient to activate them to induce antennal grooming.

We used this assay to test individual flies before bilateral antennal ablation (ctl) and at 7 dpa (Fig. 3). Flies expressing GFP in JO neurons failed to elicit antennal grooming following red-light exposure at 7 dpa (Movie S1, Fig. 3). In contrast, flies with JO-specific NMN-D expression continued to elicit antennal grooming at 7 dpa (Movie S2, Fig. 3). Remarkably, the evoked grooming behavior remained equally robust at 14 dpa (Fig. S4). The preservation of synaptic connectivity was dependent on low NMN levels, as flies expressing NMN-D^dead^ in JO neurons failed to elicit antennal grooming upon red-light exposure at 7 dpa (Movie S3, Fig. 3). Our findings demonstrate that lowering NMN potently attenuates axon death signaling, which is sufficient to preserve synaptic connectivity of severed axons and synapses for weeks after injury.

**Figure 3.**
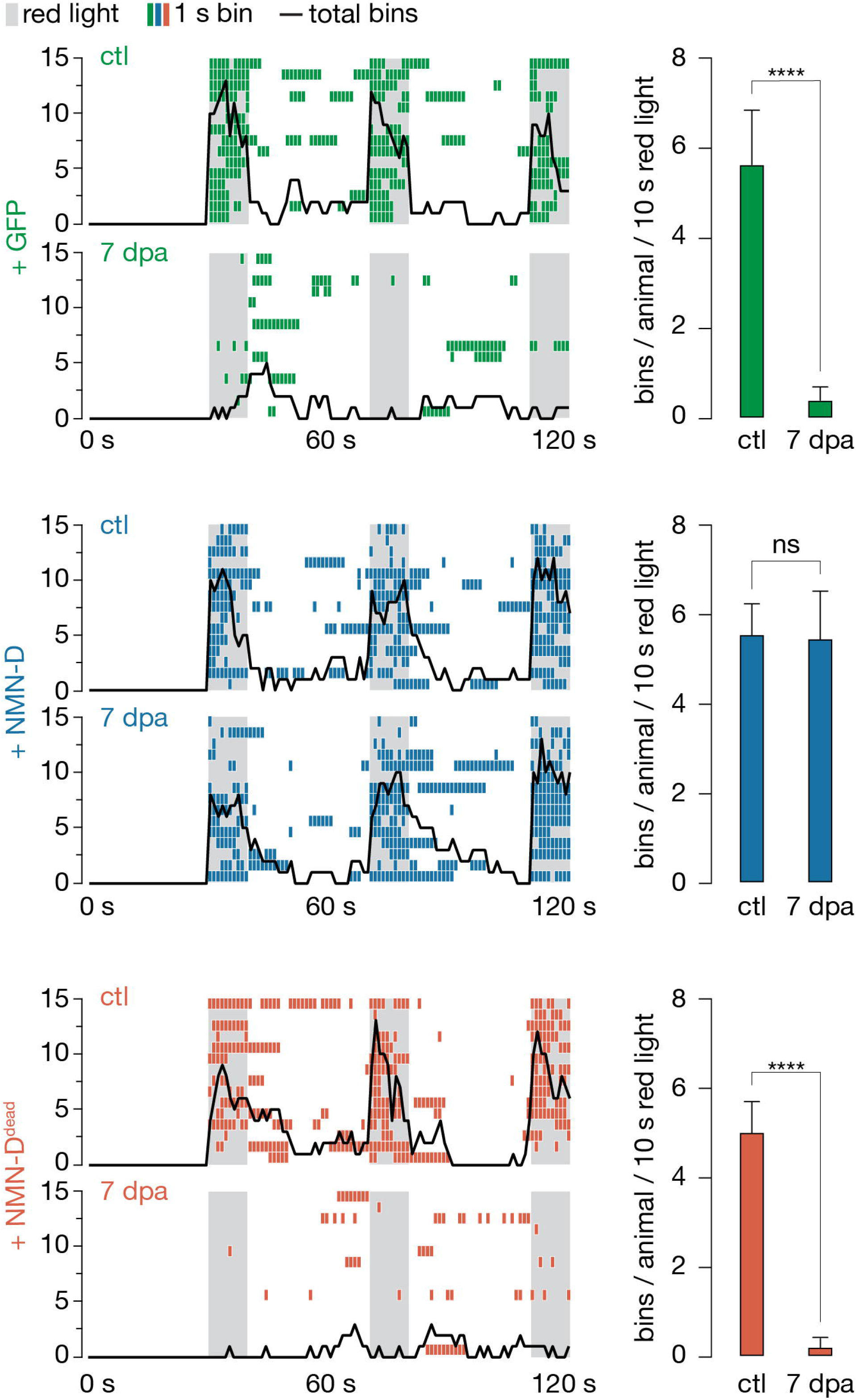
Low neuronal NMN preserves synaptic connectivity at 7 dpa. Antennal grooming induced by red light. Left: ethograms of uninjured control (ctl) and 7 dpa flies. Gray bars, 10 s red light; colored boxes, bins; black line, sum of bins (n = 15 flies). Right: average bins per animal during 10 s red-light exposure (n = 15 flies). Two-tailed t-student test, **** = *p* < 0.0001, ns = *p* > 0.05.

### mNAMPT-expressing axons degenerate faster after injury

Two enzymatic reactions synthesize NMN in mammals: NAMPT-mediated NAM and NRK1/2-mediated NR consumption. *Drosophila* lacks NAMPT activity and NMN synthesis relies solely on Nrk-mediated NR consumption (Table S2). We hypothesized that mouse NAMPT (mNAMPT) expression could increase NMN synthesis and, therefore, lead to faster injury-induced axon degeneration *in vivo* (Fig. 4A). We generated transgenic flies harboring mNAMPT under the control of UAS by targeted insertion (*attP40*). Western blots revealed readily detectable mNAMPT, suggesting proper expression of mNAMPT in fly heads (Fig. 4B).

**Figure 4.**
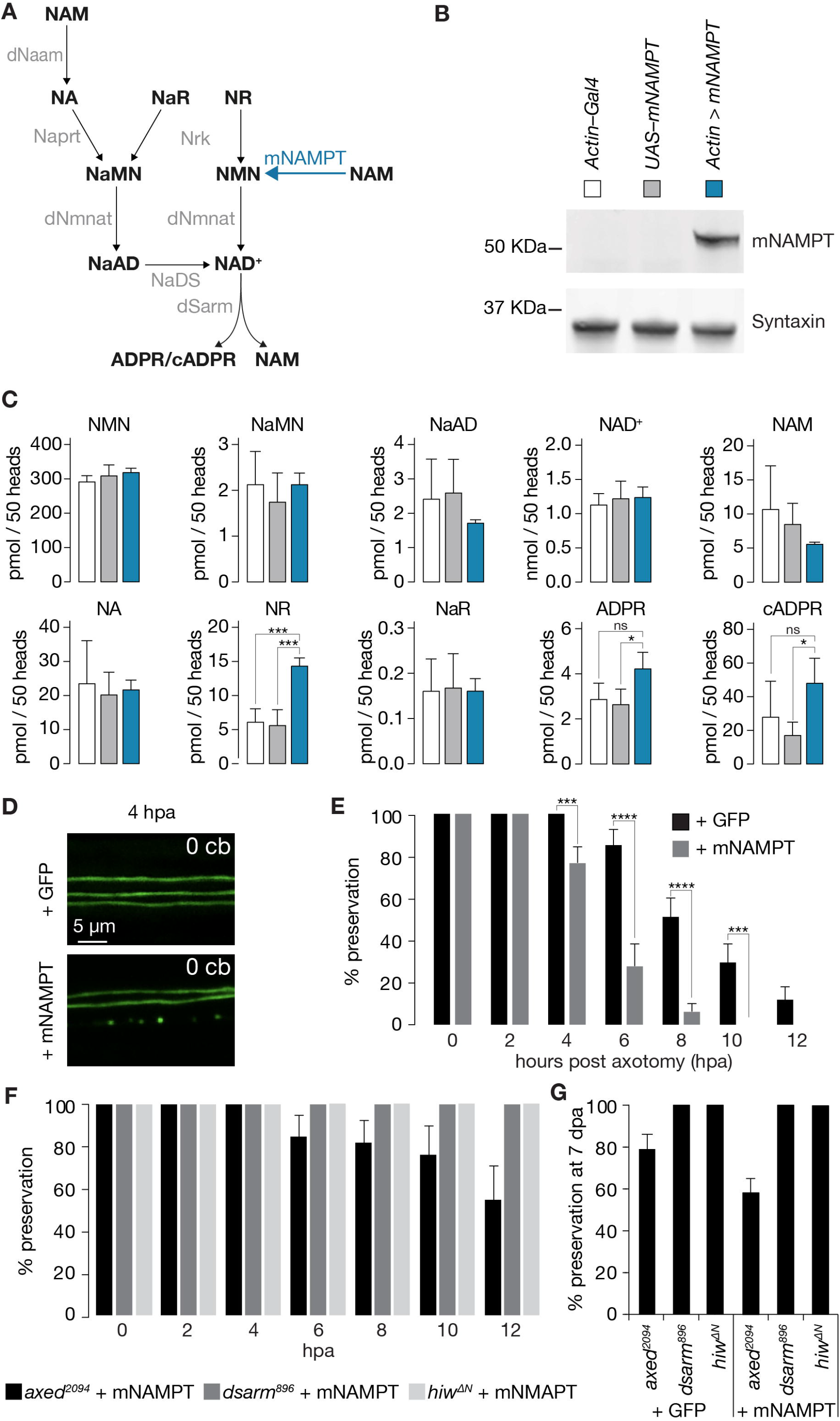
Faster injury-induced axon degeneration through mammalian NAMPT expression. **A** *Drosophila* NAD^+^ metabolic pathway. Black, metabolites; grey, enzymes; blue, mouse NAMPT (mNAMPT). **B** Detection of mNAMPT expression in heads by Western blot. **C** Subtle changes in NAD^+^ metabolic flux by mNAMPT expression in fly heads. Extracted metabolites from 50 heads, mean ± standard deviation (*n* = 4). One-way ANOVA with Tuckey’s multiple comparisons test. **D** The expression of mNAMPT results in faster axon degeneration after injury. Examples of injured axons at 4 hours post axotomy (hpa). **E** % preservation of injured axons within 12 hours post axotomy (hpa), average ± standard error of the mean (n = 20 wings). Multiple unpaired t-tests. **F** Faster axon degeneration by mNAMPT expression requires axon death genes. % preservation of injured axons within 12 hpa, average ± standard error of the mean (n = 10 wings) **G** % preservation of injured axons at 7 dpa, average ± standard error of the mean (n = 15 wings) **** = *p* < 0.0001, *** = *p* < 0.001, * = *p* < 0.01, ns = *p* > 0.05.

We then tested the effect of mNAMPT on the NAD^+^ metabolic flux *in vivo*. Surprisingly, NAM, NMN, and NAD^+^ levels remained unchanged under physiological conditions (Fig. 4C). We noticed 3-times higher NR and a trend towards higher ADPR and cADPR (Fig. 4C).

Although mNAMPT expression failed to elevate NMN under physiological conditions, we hypothesized that mNAMPT could “boost” NMN levels after injury because of the following observations made in mammals. NMNAT2 is labile and rapidly degraded in severed axons ^15^ while NAMPT persists much longer ^19^. We speculated that in severed axons in flies, dNmnat declines similarly, but not mNAMPT. In consequence, NMN would therefore accumulate. Strikingly, in our wing injury assay, while axons with GFP showed signs of degeneration starting from 6 hours post axotomy (hpa), mNAMPT expression resulted in significantly faster axon degeneration, with signs of degeneration at 4 hpa (Fig. 4D, E).

We next asked whether the faster degeneration of mNAMPT-expressing severed axons requires axon death genes. While mutations in *dsarm* and *hiw* completely blocked the degeneration of severed axons expressing mNAMPT, *axed* showed a partial preservation of 60 % at 12 hours after injury (Fig. 4F). Importantly, a similar preservation was observed at 7 dpa. Severed GFP- and mNAMPT-expressing axons were similarly preserved in *dsarm, hiw*, and *axed* mutant backgrounds (Fig. 4G), arguing that elevated NMN levels require axon death signaling to initiate the degeneration of severed axons.

Overall, our data suggest that NMN accumulation after injury triggers axon degeneration in *Drosophila* through the axon death pathway. To our knowledge, we provide the first direct *in vivo* demonstration that an additional source of NMN synthesis, by the expression of mNAMPT, accelerates injury-induced axon degeneration.

### NMN activation of dSarm NADase is required for axon degeneration *in vivo*

We have shown that low NMN attenuates injury-induced axon degeneration, while a more rapid accumulation of NMN, due to ectopic expression of mNAMPT, results in faster injury-induced axon degeneration. A cell-permeable form of NMN (CZ-48) binds to and activates SARM1 by changing its conformation ^23^. Crystal structures of the ARM domain of dSarm (dSarm^ARM^) as well as the full-length human SARM1 (hSARM1) support this observation, where NMN acts as a ligand for dSarm^ARM^ ^11,43^. NMN binding to the ARM domain requires a critical residue, lysine 193 (K193), for the induction of a conformational change in the ARM domain of dSarm/SARM1. Mutations thereof (e.g., K193R) result in a dominant-negative injury-induced axon degeneration phenotype in murine cell cultures ^11,21–23,44^.

To confirm whether NMN activates dSarm *in vitro* and *in vivo*, we generated *dsarm* constructs encoding wild-type and the K193R-equivalent K450R mutation. Crucially, an isoform we used previously, dSarm(D), fails to fully rescue the *dsarm^896^* defective axon death phenotype (Fig. S5A, B) ^30^. Therefore, among the eight distinct *dsarm* transcripts, which all contain the ARM, SAM, and TIR domains, we chose the shortest coding isoform, *dsarm*(*E*) (Fig. S6A). We generated un-tagged and C-terminal FLAG-tagged dSarm(E), with and without K450R, under the control of UAS and confirmed the FLAG-tagged proteins are expressed at similar levels in S2 cells (Fig. 5A). We also directly tested the immunopurified FLAG-tagged proteins for constitutive and NMN-inducible NADase activity (Fig. 5B). While wild-type and K450R dSarm(E) had similar constitutive activities in the presence of 25 μM NAD^+^ alone, we found that only wild-type NADase activity was induced further with the addition of 50 μM NMN. At the same time, K450R remained essentially unchanged (Fig. 5C). This confirmed the critical role of this residue in NMN-dependent triggering of dSarm NADase, similar to its role in hSARM1^11,21^.

**Figure 5.**
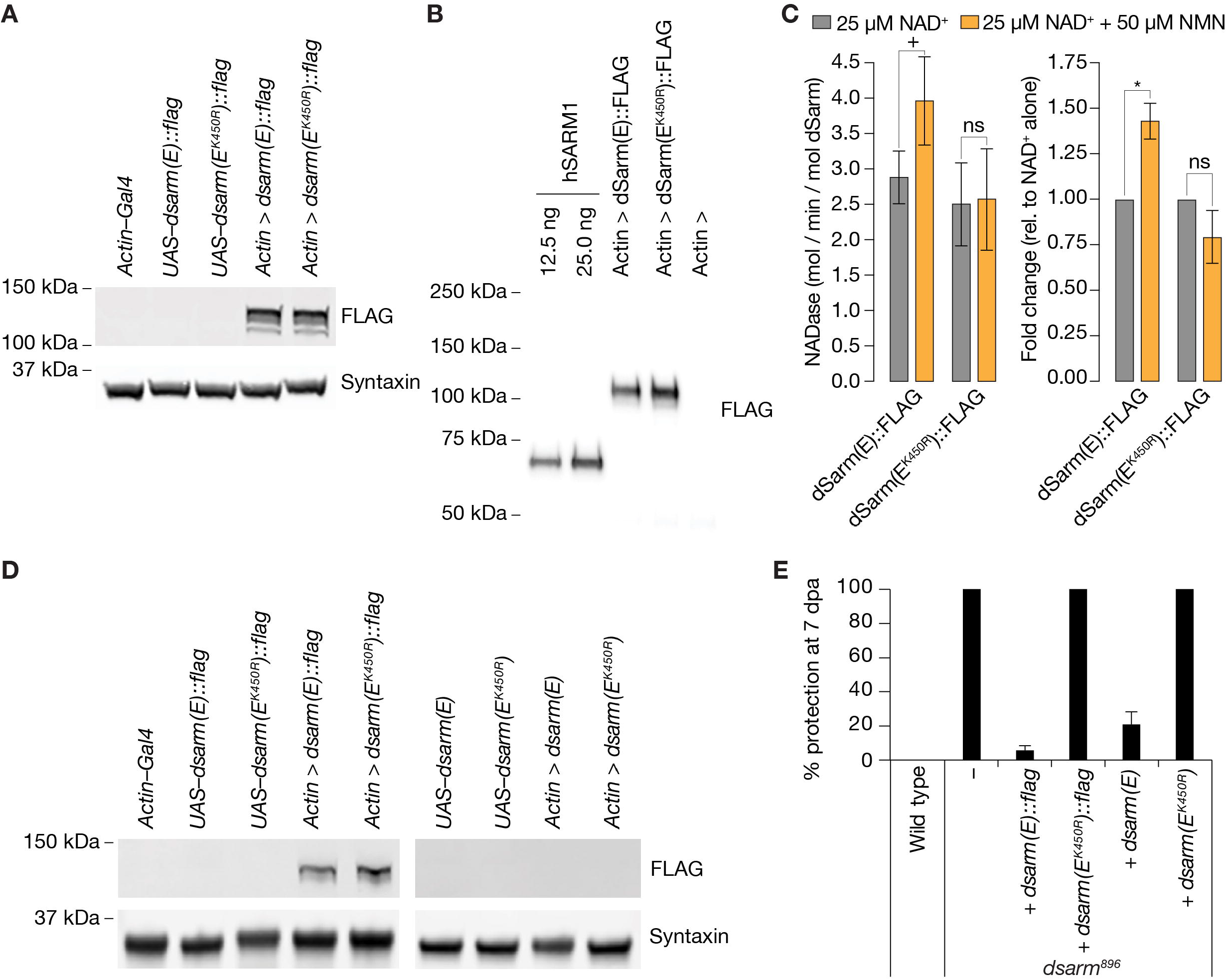
NMN inducibility of dSarm NADase is required for axon degeneration *in vivo*. **A** Expression and detection of wild-type and K450R dSarm(E) variants in S2 cells. **B** Immunoblot of immunopurified dSarm(E)::FLAG and dSarm(E^K450R^)::FLAG. Known amounts of immunopurified human SARM1 (hSARM1) were used to quantify the levels of immunopurified dSarm(E); 12.5 ng = 154.5 fmol hSARM1; 25 ng = 309 fmol hSARM1 **C** NADase activity of dSarm. Left: NADase activity (mol NAD consumed / min / mol dSarm) of immunopurified dSarm(E)::FLAG and dSarm(E^K450R^)::FLAG in the presence of 25 μM NAD^+^, or 25 μM NAD^+^ + 50 μM NMN. Right: degree of MNM induction (fold-change relative to NAD^+^ alone). Mean ± standard error of mean (*n* = 5). Control immunoprecipitations (using extracts from *Actin–Gal4* transfected S2 cells) revealed no non-specific NAD^+^-consuming activity on equivalent amounts of bead / antibody complexes compared to that used in the dSarm(E) activity assays (*n* = 5). Multiple paired t-test with false discovery rate (FDR) correction. **D** Equal expression levels of dSarm(E) variants in *Drosophila* heads. **E** Rescue experiments of dSarm(E) variants in *dsarm^896^* mutant clones. dSarm(E) rescues, while dSarm(E^K450R^) fails to rescue the *dsarm^896^* axon death defective phenotype. % preservation of severed axons at 7 dpa, average ± standard error of mean (*n* = 15 wings), ns = *p* < 0.051, + = *p < 0,051*, * = *p* < 0.05.

Next, we generated transgenic flies expressing tagged and untagged wild-type and mutant dSarm(E) variants–by targeted insertion of the *UAS–dsarm(E*) plasmids (*attP40*)- and confirmed pan-cellular expression of the FLAG-tagged variants by immunoblotting (Fig. 5D). We used all variants for *dsarm^896^* axon death defective rescue experiments in our wing injury assay. We found that expression of wild-type dSarm(E) (both tagged and untagged) almost entirely rescued *dsarm^896^* mutants, whereas dSarm(E^K450R^) proteins completely failed to rescue the phenotype at 7 dpe (Fig. 5E). We demonstrate that a noninducible NADase variant, dSarm(E^K450R^), in the absence of wild-type dSarm, fails to execute injury-induced axon degeneration *in vivo*.

### Low NMN delays neurodegeneration triggered by loss of *dnmnat*

Lowering levels of NMN confers very robust protection against axon degeneration in *Drosophila*, similar to or even more potent than that achieved by targeting other mediators of axon degeneration, such as *hiw, dsarm, axed*, and the over-expression of *dnmnat* (*dnmnat^OE^*) ^9,29,30,34–36^. We therefore assessed the genetic interaction among these regulators *in vivo*.

The current model, supported by our data, predicts that NMN accumulation occurs upstream of dSarm activation. Consistent with this, the induced expression of constitutively active dSarm lacking its inhibitory ARM domain (dSarm^ΔARM^) is sufficient to pathologically deplete NAD^+^, thereby triggering axon- and neurodegeneration in the absence of injury ^10,30^. We asked whether lowering levels of NMN can delay or prevent neurodegeneration induced by dSarm^ΔARM^-mediated NAD^+^ depletion. *dsarm^ΔARM^* clones with forced NAD^+^ depletion rapidly degenerated within 5 days after adults were born (days post eclosion, dpe) (Fig. 6A). As expected, lowering NMN levels by NMN-D in *dsarm^ΔARM^* clones did not alter the kinetics of neurodegeneration (Fig. 6A). These observations further support that NMN accumulation occurs upstream of dSarm activation, and that once neuronal NAD^+^ is low, neurodegeneration cannot be halted by low NMN.

**Figure 6.**
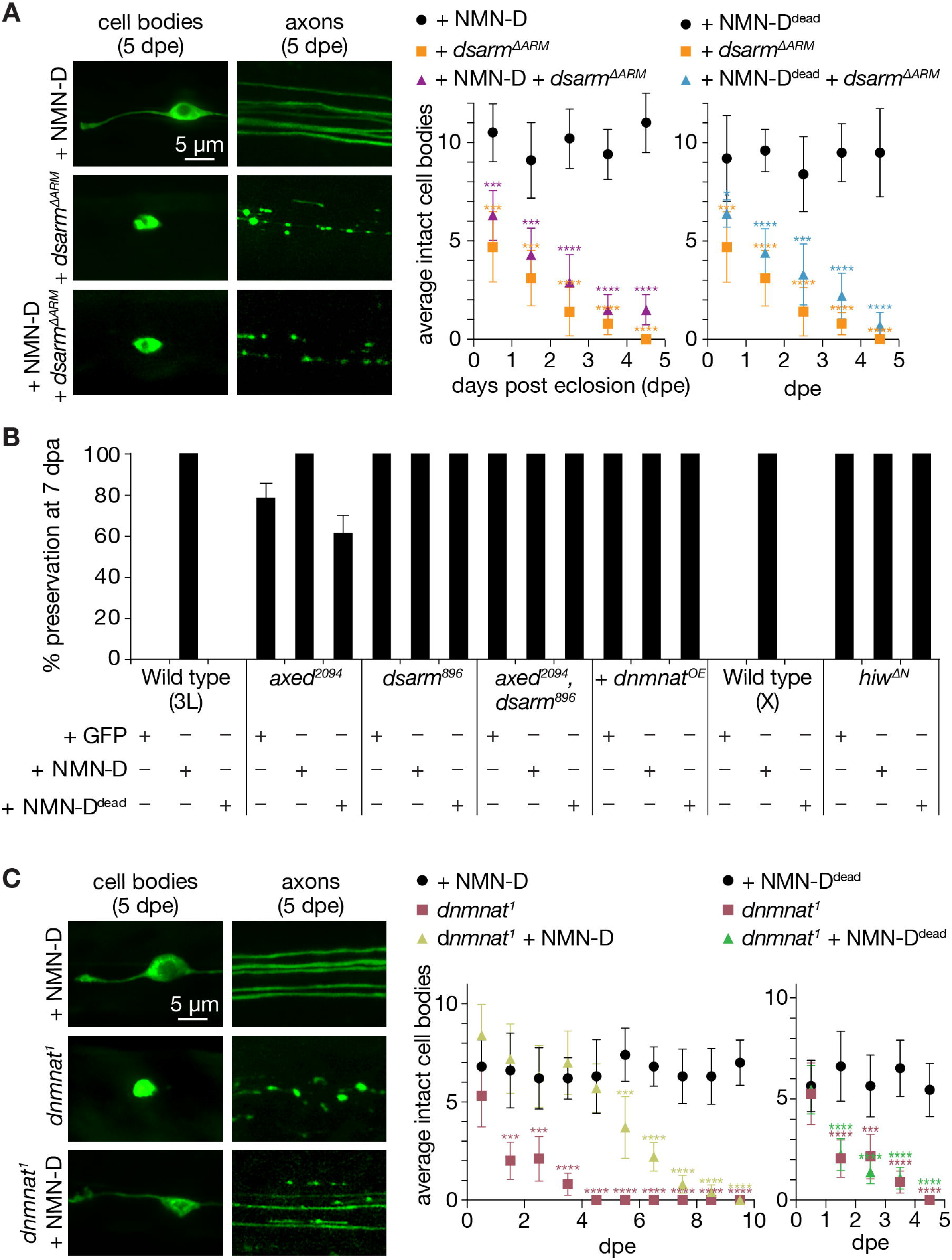
Low neuronal NMN delays neurodegeneration triggered by the loss of *dnmnat*. **A** Low NMN fails to prevent neurodegeneration triggered by dSarm^ΔARM^-mediated NAD^+^ depletion. Top: examples of cell bodies and axons at 5 days post eclosion (dpe). Bottom: quantification of intact cell bodies, average ± 95 % confidence interval (CI), (*n* = 10 wings). B Epistasis analysis of low NMN with axon death signaling genes. Low NMN does not alter *axed, dsarm, hiw*, or *dnmnat* overexpression (+ *dnmnat^OE^*) phenotypes in the wing injury assay. % preservation of injured axons at 7 dpa, average ± standard error of the mean (*n* = 15 wings). C Low NMN delays neurodegeneration triggered by the loss of *dnmnat*. Top: examples of cell bodies and axons at 5 dpe. Bottom: quantification of intact cell bodies, average ± 95 % CI, (*n* = 10 wings). Multiple unpaired t-tests. All tests are compared to the control group (e.g., black dots). **** = *p* < 0.0001, *** = *p* < 0.001, * = *p* < 0.05.

We also asked whether lowering NMN interferes with loss-of-function mutations of *axed, dsarm*, double mutants, *hiw*, and *dnmnat^OE^*. As expected, the attenuated axon death phenotype of low NMN did not change in these mutant backgrounds at 7 dpe (Fig. 6B).

Loss-of-function mutations in *dnmnat* also activate axon death signaling, leading to neurodegeneration in the absence of injury ^30^. dNmnat is the sole enzyme with NAD^+^ biosynthetic activity in *Drosophila; dnmnat^1^* mutant clones lack NMN consumption and NAD^+^ synthesis ^45^, and they degenerate with similar kinetics as *dsarm^ΔARM^* clones ^30^. We asked whether low NMN levels can delay or prevent *dnmnat^1^*-induced neurodegeneration. Surprisingly, although not expected to restore NAD^+^ synthesis, lowering NMN levels resulted in a significant delay of neurodegeneration (Fig. 6C). Between 1–5 dpe, while clones with *dnmnat^1^* fully degenerated, NMN-D expressing *dnmnat^1^* clones remained morphologically intact, similar to controls. After 5 days, these clones gradually started to deteriorate (Fig. 6C). This protective delay of neurodegeneration depends on lowering NMN levels, as the expression of NMN-D^dead^ completely failed to protect *dnmnat^1^* neurons (Fig.6C).

To confirm that NMN-D delays *dnmnat^1^*-mediated neurodegeneration, we wanted to generate tissue-specific CRISPR/Cas9 *dnmnat* knockouts. CRISPR/Cas9 tools significantly facilitate genetics in *Drosophila*. Rather than using mutants in a specific genomic locus, mutations are induced by co-expressing Cas9 and *sgRNAs*.

First, we generated transgenic flies harboring *tRNA*-flanked *sgRNAs* to target four distinct loci in *dnmnat* under the control of UAS ^46^. We made similar transgenic flies to target all other axon death genes (Fig S6A). We then tested our novel tools for their ability to attenuate axon death by assessing preserved axonal morphology (Fig. S6B) and synaptic connectivity after injury (Fig. S6C). We found that the preservation was dependent on the combination of the tissue (e.g., Gal4 driver) and the Cas9 source (e.g., *UAS–cas9* vs. *actin–cas9*). Our observations highlight that the combination of Cas9 and *sgRNAs* has to be carefully determined in each tissue targeted by CRISPR/Cas9.

We then asked whether *dnmnat^sgRNAs^* can trigger neurodegeneration by analyzing neuronal survival (Fig. S5D) and synaptic connectivity over time (Fig. S6E). Notably, we observed synthetic lethality in *UAS-dnmnar^sgRNAs^ actin–Cas9* flies; we thus used *UAS–cas9*. Neurons with CRISPR/Cas9-targeted *dnmnat* degenerated as fast as *dnmnat^1^* mutants (Fig. S5D). In line with these findings, we observed reduced synaptic connectivity in 7- and 14-day old flies (Fig. S6E). We also found the expression of NMN-D in *dnmnat^sgRNAs^* clones resulted in similar neuroprotection as observed with *dnmnat^1^* mutants. They remained morphologically intact during the first 5 days and then gradually degenerated (Fig. S6F). Therefore, NMN-D can also delay neurodegeneration in CRISPR/Cas9-targeted *dnmnat* clones by preventing NMN accumulation.

Taken together, our *in vivo* results suggest that in the absence of *dnmnat*, NMN-D prevents NMN accumulation and therefore delays neurodegeneration. However, neurons subsequently degenerate because NAD^+^ synthesis halts and NAD^+^ gradually decays below the threshold of survival. Similarly, NMN-D fails to delay neurodegeneration when dSarm^ΔARM^ forcefully depletes NAD^+^. These results support the role of NMN as an activating rather than an executing function in axon death signaling.

## Discussion

Here we investigate how lowering of the NAD^+^ precursor metabolite NMN influences axon survival in *Drosophila*, using robust expression of prokaryotic NMN-D, as demonstrated with newly generated PncC antibodies. When expressed, NMN-D consumes NMN to synthesize NaMN in *Drosophila* heads. While the protection by NMN-D could in principle reflect an inhibitory effect of NaMN ^47^, the additional acceleration of degeneration by mNAMPT strongly argues that NMN is a key mediator of dSarm-driven axon degeneration. In the context of injury-induced axon degeneration, neuronal expression of NMN-D to keep NMN low is sufficient to block axon death signaling: severed axons with NMN-D remain morphologically preserved for the lifespan of flies, and circuit-integrated for weeks after injury. Neurodegeneration induced by dNmnat depletion is also delayed by low NMN levels. Our data indicate that NMN is a key mediator of axon degeneration in *Drosophila*, acting as an activator of dSarm in the axon death pathway *in vivo*. This is consistent with observations in mammals ^19,22,23^, and with previously reported direct binding of NMN to the dSarm ARM domain^11^.

The discovery and characterization of the axon death signaling pathway revealed four major players mediating axonal degeneration in *Drosophila*. Loss-of-function mutations in *hiw, dsarm* and *axed*, as well as *dnmnat* over-expression robustly inhibit injury-induced axon degeneration ^9,29,30,34,35^. We now show that lowering NMN levels has an equally potent protective effect, adding NMN as an additional mediator to the signaling pathway. Synaptic connectivity of severed axons is also preserved for weeks, comparable to, or even better than that accomplished by *hiw, dsarm* and *axed* mutants ^30^, and *dnmnat* over-expression ^36^.

Our demonstration of NMN as a mediator of axon degeneration in *Drosophila* addresses an important question in the field. While recent discoveries are confirming the original finding of a pro-degenerative action of NMN ^19^, the role of NMN in axon degeneration has also been questioned, especially in *Drosophila*. Given the absence of NAMPT in flies, it is tempting to speculate that NMN–as a minor intermediate of the NAD^+^ metabolic pathway–is not a primary contributor in injury-induced axon degeneration ^32^. However, we provide compelling evidence that NMN is not only present in flies, as previously reported ^48^, but that its accumulation causes axon degeneration. In line with other studies, we show further proof that NMN acts as an activator of dSarm *in vitro* by using a new *dsarm* isoform, *dsarm(E)*, which is fully functional in axon death signaling. Its non-inducible variant, dSarm(E^K450R^) fails to rescue the attenuated axon degeneration phenotype in neurons lacking *dsarm*. Together with the reported dominant negative effect of SARM1(K193R) in mice our observations further support that NMN activation of dSarm also occurs *in vivo*.

The previously published NMNd revealed partially protected axons 7 days after injury in the wing ^33^, while our NMN-D extends protection to 50 days. This difference is likely due to the N-terminal GFP tag in GFP::NMN-D, which can increase protein stability ^37^. This is supported by our newly generated PncC antibodies and suggests that NMN-D expression levels dictate reduction of NMN, and therefore protection of severed axons.

We also demonstrate that increasing the synthesis of NMN provokes a faster degeneration of severed axons *in vivo*, which requires all axon death mediators. Mammals synthesize NMN with two distinct enzymatic reactions: NR consumption by NRK1/2 and NAM consumption by NAMPT, both ensuring NMN supply. In *Drosophila*, NAMPT activity is absent, NMN appears to be synthesized by Nrk alone, yet dietary supplementation might also contribute ^49^. We used mouse NAMPT as an extra source of NMN synthesis. However, in contrast to the NMN-D induced change in the NAD^+^ metabolic flux, mNAMPT had only a minor impact under physiological conditions. It is challenging to measure the specific axonal rise in NMN after injury *in vivo*. However, NMNAT2 is rapidly disappearing in served axons ^15^, and so is dNmnat in *Drosophila* axons and synapses ^29^, through PHR1 and Hiw, respectively, while NAMPT persists longer^19^. It is therefore likely that persisting mNAMPT in severed *Drosophila* axons continues NMN synthesis, leading to faster NMN accumulation, dSarm activation, and faster axon degeneration.

Finally, we expanded our investigations beyond injury, by looking at NMN in a model of neurodegeneration. dNmnat is essential for NAD^+^ synthesis. While neuronal clones with mutant *dnmnat^1^* start to degenerate after they are born, intriguingly, the co-expression of NMN-D resulted in a preservation of neuronal morphology for at least 5 days, before degeneration started with similar kinetics. Our results suggest that a rise in NMN is a trigger for neurons to degenerate also in this model, rather than the lack of NAD^+^ biosynthesis, at least within the first 5 days. Once NAD^+^ levels drop beyond neuronal survival, neurons eventually degenerate. This is supported by observation with forced NAD^+^ depletion by dSarm^ΔARM^ ^30^, and by the inhibition of NAD^+^ biosynthesis in murine neurons with FK866 ^19^. Still, it is surprising to observe that neurons lacking NAD^+^ synthesis are capable of surviving for days. It suggests either that the NAD^+^ turnover is slower than expected ^50^ or mechanism are in place to compensate for NAD^+^ loss, at least in the short term.

In conclusion, NMN is a potent mediator of axon- and neurodegeneration in *Drosophila*. Our newly developed NMN-D tool will be useful in many degenerative aspects beyond injury, such as in axon morphogenesis and maintenance ^51^, and also in dendrite pruning ^52^. Our metabolic analyses further demonstrate that *Drosophila* serves as an excellent model system to study NAD^+^ metabolism *in vivo*.

## Supplementary Figure Legends

**Figure S1. Partial protection of severed axons at 7 dpa by previously published NMN-Deamidase (NMNd). A** Control and 7 days post axotomy (7 dpa) examples of GFP- and NMNd-expressing axons in the wing injury assay. Cell bodies in the cut-off distal wing are immediately counted to determine severed axons. After injury, the number of neuronal cell bodies (cb) that remain attached in the proximal wing–displayed in the upper right corner of each example–indicates how many uninjured, thus expected axons remain in the nerve bundle. **B** Axon death quantification. Left: average scores of uninjured control, debris, and severed intact axons (white, gray, and black, respectively; *n* = 15 wings). Right: % preservation of injured axons at 7 dpa, average ± standard error of the mean (*n* = 15 wings).

**Figure S2. Increased NMN-D detected by anti-PncC antibodies. A** NMN-D expression and detection by anti-PncC antibodies in S2 cells. B Increased levels of NMN-D compared to NMNd in heads of *Drosophila*. Arrow, predicted NMN-Deamidase; arrowhead, potential degradation product. **C** Quantification of Western blot PncC-immunoreactivity by densitometry. 2 heads / lane; mean ± standard deviation (n = 4); a.u., arbitrary units. Oneway ANOVA with Tuckey’s multiple comparisons test. *** =*p* < 0.001, ns, not significant, = *p* > 0.05.

**Figure S3. Quantification of axonal phenotypes.** Quantification of uninjured control (white), debris (grey), and severed intact (black) axons in uninjured and injured wings (n = 15; 7 dpa – and +, respectively).

**Figure S4. Low neuronal NMN preserves synaptic connectivity for weeks after injury.** Antennal grooming induced by red light. Left: ethograms of uninjured control (ctl), 7 and 14 dpa flies. Gray, 10 s red light; blue boxes; bins; black line, sum of bins (n = 15). Right: average bins per animal during 10 s red-light exposure (n = 15 flies). Two-tailed t-student test, ** = *p* < 0.01, ns = *p* > 0.05.

**Figure S5. Partial rescue of *dsarm^896^* axon death defective phenotype by dSarm(D) isoform. A** dSarm(D)::GFP is not detected by GFP immunoreactivity in Western blots of fly heads. B Expression of dSarm(D) and dSarm(D)::GFP partially rescues the *dsarm^896^* axon death defective phenotype. % preservation of injured axons at 7 dpa, average ± standard error of the mean (*n* = 15 wings).

**Figure S6. Newly generated axon death gene CRISPR/Cas9 *sgRNAs*. A** Targets and orientation of axon death gene *sgRNAs*. Schematic genomic loci and indicated isoforms. Black arrows, *sgRNA* target and orientation. **B** Preservation of severed axons by attenuated axon death CRISPR/Cas9 tools depends on Cas9 source. % preservation of severed axons (average ± 95 % CI) at 7 dpa (n = 15 wings). **C** Preservation of synaptic connectivity after injury by attenuated axon death CRISPR/Cas9 tools depends on Cas9 source. Quantification of bins, average ± standard deviation (*n* = 15 flies). Two-way ANOVA with Tuckey’s multiple comparisons test. D Activation of axon death by targeting *dnmnat* with CRISPR/Cas9 phenocopies *dnmnat^1^* mutants (*dnmnat^sgRNAs^ + UAS-cas9*). Quantification of intact cell bodies, average ± 95 % CI (n = 10 wings). Multiple unpaired t-tests. All tests are compared to the control group (e.g., black dots). E Activation of axon death by targeting *dnmnat* with CRISPR/Cas9 reduces evoked grooming behavior over time (*dnmnat^sgRNAs^ + UAS–cas9*). Quantification of bins, average ± standard deviation (*n* = 15 flies). Two-way ANOVA with Tuckey’s multiple comparisons test. F Low NMN delays degeneration in CRISPR/Cas9-targeted *dnmnat* neurons (*dnmnat^sgRNAs^ + UAS–cas9*). Quantification of intact cell bodies, average ± 95 % CI (n = 10 wings). Multiple unpaired t-tests. All tests are compared to the control group (e.g., black dots). **** = *p* > 0.0001, *** = *p* < 0.001, ** = *p* > 0.01, * = *p* < 0.05.

## Supplementary Table Legends

**Table S1. Genotypes in each display item.** Abbreviations: *mCD8::GFP = UAS-mCD8::GFP, dpn = dpr1–Gal4*

**Table S2. Detected activities of NAD^+^ metabolic pathway enzymes in extracts of *Drosophila* heads.** Enzymes are shown in Figure 2A, Qaprt: quinolinate phosphoribosyltransferase, catalyzes quinolinate to NaMN from the de novo NAD^+^ synthesis pathway. Values are listed as mean ± standard deviation (n = 2); nd, not detected.

## Supplementary Movie Legends

**Movie S1. Related to Figure 3.**

Examples of red light-stimulated wild-type flies expressing CsChrimson and GFP in JO neurons, uninjured and at 7 dpa.

**Movie S2. Related to Figure 3.**

Examples of red light-stimulated wild-type flies expressing CsChrimson and GFP::NMN-D in JO neurons, uninjured and at 7 dpa.

**Movie S3. Related to Figure 3.**

Examples of red light-stimulated wild-type flies expressing CsChrimson and GFP::NMN-D^dead^ in JO neurons, uninjured and at 7 dpa.

## Author contributions

ALR, AL and LJN conceived the study and designed experiments. CA, MG, LC, NR and GO performed enzymatic assays. AL generated NMN-D plasmids and transgenic NMN-D flies. ALR performed Western blots, microscopy, cell cultures, and IPs. ALR and MP performed optogenetics. MK generated mNAMPT plasmid and transgenic mNAMPT flies. ALR and LJN generated CRISPR/Cas9 *sgRNA* plasmids and transgenic *sgRNA* flies. JG performed NADase assays. ALR and MK collected heads for metabolomics. ALR generated *dsarm(E)* plasmids and transgenic *dsarm(E)* flies. ALR, JG, GO, MPC, AL and LJN wrote the manuscript.

## Acknowledgments

We thank Dr. Jemeen Sreedharan for support in generating transgenic flies, and Dr. Julijana Ivanisevic and Dr. Hector Gallart-Ayala from the Metabolomics Unit at University of Lausanne for metabolic analyses. This work was supported by a UK Biotechnology and Biological Sciences Research Council (BBSRC) / AstraZeneca Industrial Partnership award (BB/S009582/1a) to JG; funds from the Italian Grant RSA 2018-20 from UNIVPM to GO; a Sir Henry Wellcome Postdoctoral Fellowship from the Wellcome Trust (210904/Z/18/Z) to AL; the John and Lucille van Geest Foundation to MPC; and a Swiss National Science Foundation SNSF Assistant Professor award (176855), the International Foundation for Research in Paraplegia (P180), and SNSF Spark (190919) to LJN.

## Materials & Methods

### Fly genetics

Flies (*Drosophila melanogaster*) were kept on Nutri-Fly Bloomington Formulation (see resources table) with dry yeast at 20 °C unless stated otherwise. The following genders were scored as progeny from MARCM crosses: females (X chromosome); and males & females (autosomes, chromosomes 2L, 2R, 3L, and 3R). We did not observe any gender-specific differences in clone numbers or axon death phenotype. Gender and genotypes are listed in Table S1.

### NAD-related enzymes assay

#### Sample extraction

Fly heads (previously collected and frozen, 50 weighed heads per sample) were ground in liquid N_2_ and sonicated after resuspension in 200-250 μl of 50 mM Tris-HCl pH 7.5, 0.3 M NaCl, 1 mM PMSF, and 2 μg/ml each of aprotinin, leupeptin, chimostatin, pepstatin and antipain. The suspension was centrifuged at 40000 g for 20 min at 4 °C. The supernatant was passed through a G-25 column (GE Healthcare) equilibrated with 50 mM Tris-HCl pH 7.5, 0.3 M NaCl to remove low molecular weight compounds that interfere with the enzymatic assays. Protein contents were measured with the Bio-Rad Protein Assay.

#### Nampt, Nrk, Naprt, and Qaprt activities

Enzymes were assayed according to ^39^ with minor modifications. Briefly, their formed reaction products, either NMN or NaMN, were converted to NAD using ancillary enzymes PncC (bacterial NMN deamidase), NadD (bacterial NaMN adenylyltransferase), and NadE (bacterial NAD synthase), followed by quantification of NAD with a fluorometric cycling assay ^39^.

First, mononucleotide products were converted to NaAD in dedicated assay mixtures as described below.

##### Nampt

The assay mixture consisted of ethanol buffer (30 mM HEPES/KOH pH 8.0, 1 % v/v ethanol, 8.4 mg/ml semicarbazide), 40 mM HEPES/KOH pH 7.5, 10 mM KF, 10 mM MgCl_2_ 2.5 mM ATP, 0.3 mM NAM, 2 mM PRPP, 6 U/ml ADH, 0.067 mg/ml BSA, 1 U/ml NadD, 0.03 U/ml PncC, in a final volume of 100 μl.

##### Nrk

The assay mixture was similar to the one of NAMPT, lacking PRPP, with 2 mM NR instead of NAM, and with 5 μM FK866.

##### Naprt

The assay mixture included ethanol buffer, 40 mM HEPES/KOH pH 7.5, 10 mM KF, 20 mM MgCl_2_ 2.5 mM ATP, 2 mM PRPP, 0.5 mM NA, 6 U/ml ADH, 0.067 mg/ml BSA and 1 U/m NadD.

##### QaPRT

The assay mixture included ethanol buffer, 30 mM potassium phosphate buffer pH 7.0, 10 mM KF, 5 mM MgCl_2_, 2.5 mM ATP, 2 mM PRPP, 0.3 mM QA, 6 U/ml ADH, 0.067 mg/ml BSA and 1 U/ml NadD.

Second, aliquots of the assay mixtures were withdrawn at different incubation times at 37 °C, treated with perchloric acid to stop the reactions, and incubated in a NadE mixture to transform NaAD into NAD. Third, NAD was quantified with fluorometric cycling ^39^.

#### dNaam activity

NA was converted to NaAD by the consecutive actions of the ancillary enzymes PncB (bacterial NaPRT) and NadD. The reaction mixture consisted of ethanol buffer, 40 mM HEPES/KOH pH 7.5, 10 mM KF, 10 mM MgCl_2_ 2.5 mM ATP, 2 mM PRPP, 0.3 mM NAM, 6 U/ml ADH, 0.067 mg/ml BSA, 0.5 U/ml PncB and 1 U/ml NadD. A control mixture was prepared in the absence of NAM. The generated NaAD was converted to NAD which was quantified as described above.

#### dNmnat and NaDS activities

Enzymatic activities were determined by directly measuring the newly synthesized NAD as follows:

##### dNmnat

The assay mixture consisted of 40 mM HEPES/KOH pH 7.5, 10 mM KF, 1 mM DTT, 25 mM MgCl_2_,1 mM ATP, 1 mM NMN. NMN was omitted in a control mixture.

##### NaDS

the assay mixture included 50 mM HEPES/KOH pH 7.5, 10 mM KF, 50 mM KCl, 5 mM MgCl_2_, 4 mM ATP, 20 mM glutamine, 1 mM NaAD. NaAD was omitted in a control mixture. The dNmnat and NaDS assay mixtures were incubated at 37 °C, aliquots were withdrawn and immediately subjected to acidic treatment to stop the reaction at various times. The newly synthesized NAD was quantified as described above.

One Unit (U) above refers to the amount of enzyme that forms 1 μmol/min of product at the indicated temperature, under conditions of initial velocity, e.g., less than 20 % of substrate consumption. Other activity values are reported as pmol/hour/50 heads of product formed and are means ± standard deviation of two independent experiments. The ancillary bacterial enzymes PncC, NadD, and NadE were prepared as described ^39^, whereas *Staphylococcus aureus* PncB was prepared according to ^38^.

### NAD^+^ metabolite quantification by *LC-MS/MS*

#### Sample extraction

Fly heads (50 per sample) were extracted with 125 μl of ice-cold methanol containing stable isotope-labeled (e.g., internal standard or ISTD) metabolites. Sample extracts were vortexed and centrifuged (15 min, 14000 rpm at 4 °C). The resulting supernatant was collected and evaporated to dryness in a vacuum concentrator (LabConco, Missouri, US). Sample extracts were reconstituted in 50 μl of ddH_2_o prior to LC-MS/MS analysis.

#### LC-MS/MS

Extracted samples were analyzed by Liquid Chromatography coupled with tandem mass spectrometry (LC-MS/MS) in positive electrospray ionization (ESI) mode. An Agilent 1290 Infinite (Agilent Technologies, Santa Clara, California, US) ultra-high performance liquid chromatography (UHPLC) system was interfaced with Agilent 6495 LC-MS QqQ system equipped with an Agilent Jet Stream ESI source. This LC-MS/MS was used to quantify the intermediates implicated in NAD^+^ *de novo synthesis* and *salvage pathways* ^40^.

The separation of NAD^+^ metabolites implicated in salvage and Preiss-Handler pathway was carried out using the Scherzo SMC18 (3 μm 2.0 mm x 150 mm) column (Imtakt, MZ-Analysentechnik, Mainz, Germany). The two mobile phases were composed of 20 mM ammonium formate and 0.1 % formic acid in ddH_2_0 (= A) and acetonitrile: ammonium formate 20 mM and 0.1 % formic acid (90:10, v/v) (= B). The gradient elution started at 100 % A (0 - 2 min), reaching 100 % B (2 - 12 min), then 100% B was held for 3 min and decreased to 100 % A in 1 min following for an isocratic step at the initial conditions (16 - 22 min). The flow rate was 200 μl/min, the column temperature 30 °C and the sample injection volume 2 μl. To avoid sample carry-over, the injection path was cleaned after each injection using a strong solvent (0.2 % formic acid in methanol) followed by a weak solvent (0.2 % formic acid in ddH_2_0).

AJS ESI source conditions operating in positive mode were set as follows: dry gas temperature 290 °C, nebulizer 45 psi and flow 12 l/min, sheath gas temperature 350 °C and flow 12 l/min, nozzle voltage +500 V, and capillary voltage +4000 V. Dynamic Multiple Reaction Monitoring (DMRM) acquisition mode with a total cycle of 600 ms was used operating at the optimal collision energy for each metabolite transition.

#### Data processing

Data was processed using Mass Hunter Quantitative (Agilent). For absolute quantification, the calibration curve and the internal standard spike were used to determine the response factor. Linearity of the standard curves was evaluated using a 14-point range; in addition, peak area integration was manually curated and corrected where necessary. Concentration of metabolites were corrected for the ratio of peak area between the analyte and the ISTD, to account for matrix effects.

### S2 cell culture

*Drosophila* Schneider cells (S2, Thermo Fisher) were maintained at 26 °C in *Drosophila* Schneider’s medium (Thermo Fisher) supplemented with 10 % Fetal Bovine Serum (Thermo Fisher) and 1xPenicillin-Streptomycin (Thermo Fisher). 8 x 10^5^ cells were plated out in 10 mm plates 24 h prior to transfection. Cells were co-transfected with either *UAS–dsarm(E)::flag, UAS–dsarm(E^K193R^)::flag, UAS–GFP::NMN-Deamidase, or UAS–GFP::NMN-Deamidase^dead^* constructs and *pAc–GAL4* (Addgene) to a final concentration of 10 μg DNA/well using Mirus TransIT-Insect (Mirus Bio). 48 h post-transfection, cells were harvested with the original medium in tubes on ice. Cells were centrifuged at 5000 g for 30 s, the supernatant discarded, the cells resuspended in 5 ml of cold PBS and centrifuged again at 5000 g for 30 s. After discarding the supernatant, cells were resuspended in 300 μl/plate of cold KHM lysis buffer (110 mM CH_3_CO_2_K, 20 mM HEPES pH 7.4, 2 mM MgCl_2_ 0.1 mM digitonin, Complete inhibitor EDTA free (Roche)) and incubated for 10 min at 4 °C while briefly vortexing for 5 s and up-and-down pipetting 5 times every minute. Samples were then centrifuged at 3000 rpm for 5 min to pellet cell debris. Protein concentration of the supernatant was determined using the BCA protein quantification assay (ThermoFischer).

### Western blot

#### Sample preparation

##### Fly heads

whole fly heads were lysed in Laemmli buffer (2 heads/10 μl) and 10 μl loaded per well.

##### S2 cells

Protein concentration was determined as descried above, and 20 ug of protein prepared in Laemmli buffer and loaded per well.

#### Sample run

4-12 % surePAGE™ gels (genescript) were used with MOPS running buffer (for higher molecular weight proteins) or MES running buffer (for lower molecular weight proteins). Gels were subjected to 200 V. A molecular weight marker Precision Plus Protein™ Kaleidoscope™ Prestained Protein Ladder was used (Biorad). Proteins were transferred to PVDF membranes with the eBlot^®^ L1 system using eBlot^®^ L1 Transfer Stack supports (Genscript) and the resulting membranes were washed three times with TBS-T (Tris-buffered saline containing 0.1 % Tween^®^ 20 (Merk)). Membranes were blocked with 5 % milk (Carl-Roth) in TBS-T at room temperature (RT)for 1 h. Membranes were incubated at 4 °C with corresponding primary antibodies overnight (O/N). Membranes were washed three times with TBS-T for 10 min and incubated with secondary antibodies in 5 % milk in TBS-T at RT during 1 h. Membranes were washed three times with TBS-T for 10 min.

#### Antibody concentrations

##### Primary antibodies

1:5000 rabbit anti-GFP (Abcam), 1:15000 mouse anti-Tubulin (Sigma), 1:5000 rabbit anti-Tubulin (Abcam), 1:2000 rabbit anti-PncC (LubioScience), 1:1000 mouse anti-FLAG (Sigma), 1:500 mouse anti-Syntaxin (DSHB), 1:2000 anti-NAMPT (Merk).

##### Secondary antibodies

1:10000 goat anti-rabbit IgG (H+L) Dylight 800, 1:10000 goat antimouse IgG (H+L) Dylight 800, 1:10000 goat anti-rabbit IgG (H+L) Dylight 680, 1:10000 goat anti-mouse IgG (H+L) Dylight 680 (ThermoFisher), 1:10000 goat anti-rat IgG (H+L) Dylight 800 (ThermoFisher).

Signal acquisition: Fluorescent signals were acquired using Odissey® DLx (LI-COR). Images were quantified by densitometric analysis using ImageJ (NIH).

### Injury (axotomy) assays

#### Wing injury

Animals were kept at 20 °C for 5-7 days prior axotomy, unless stated otherwise. Axotomy was performed using a modification of a previously described protocol ^36^. One wing per anesthetized fly was cut approximately in the middle. The distal, cut-off part of the wing was mounted in Halocarbon Oil 27 on a microscopy slide, covered with a coverslip, and immediately used to count the amount of cut-off cell bodies (as readout for the number of injured axons) under an epifluorescence microscope. Flies were returned to individual vials. At 0, 2, 4, 6, 8, 10 and 12 h post axotomy (hpa), or 7 days post axotomy (dpa), wings were mounted onto a slide, and imaged with a spinning disk microscope to assess for intact or degenerated axons, as well as the remaining uninjured control axons.

#### Antennal ablation

Animals were kept at 20 °C for 5-7 days prior antennal ablation. Adults were aged at 20 °C for 5-7 days before performing antennal ablation^36^. Unilateral antennal ablation (e.g., removal of one antenna) was performed using high precision and ultra-fine tweezers, and flies returned to vial for the appropriate time. The ablation of 3^rd^ antennal segments did not damage the rest of the head or lead to fly mortality. At corresponding time points, adult brain dissections were performed as described ^36^: decapitated heads were fixed in 4 % formaldehyde in PTX (0.5 % Triton X-100 in PBS) for 20 min, and washed 3 x 10 min with PTX. Brain dissections were performed in PTX, and dissected brains were fixed in 4 % formaldehyde in PTX for 10 min, followed by 1 h of blocking in 10 % normal goat serum (Jackson Immuno) in PTX and an O/N incubation with the following primary antibodies at 4 °C in blocking solution: 1:500 chicken anti-GFP (Rockland), and 1:150 mouse anti-nc82 (DSHB). Brains were then washed 3x 10 min with PTX at RT, and incubated with secondary antibodies in PTX at RT for 2 h: 1:200 Dylight 488 goat anti-chicken (Jackson lab), and 1:200 AlexaFluor 546 goat anti-mouse (ThermoFisher). Brains were washed 3 x 10 min with PTX at RT, and mounted in Vectashield for microscopy.

### Time course of degenerating neurons

Wings of aged flies (0 – 10 days post eclosion (dpe)) were observed and imaged with a spinning disk microscope to assess for intact or degenerated neurons and axons.

### Transgenesis

The plasmids listed below were generated and used for PhiC31 integrase-mediated targeted transgenesis (Bestgene) (5xUAS, w^+^ marker). *attP40* target site: *UAS–GFP::NMN-Deamidase, UAS–GFP::NMN-Deamidase^dead^, UAS–dsarm(E), UAS–dsarm(E)::flag, UAS–dsarm(E^K450R^), UAS–dsarm(E^K450R^)::flag. VK37* target site: *UAS–4x(tRNA::axed^sgRNAs^), UAS–4x(tRNA::hiw^sgRNAs^), UAS–4x(tRNA::dsarm^sgRNAs^), UAS–4x(tRNA::dnmnat^sgRNAs^*).

### Optogenetics

Crosses were performed on standard cornmeal agar containing 200 μM all-trans retinoic acid in aluminum-wrapped vials to keep the progeny in the dark ^36^. CsChrimson experiments were performed in the dark, and flies were visualized for recording using an 850 nm infrared light source at 2 mW/cm^2^ intensity (Mightex, Toronto, CA). For CsChrimson activation, 656 nm red light at 6 mW/cm^2^ intensity (Mightex) was used. Red light stimulus parameters were delivered using a NIDAQ board controlled through Bonsai (https://open-ephys.org/). Red-light stimulation (10 Hz for 10 s) was followed by a 30 s interstimulus recovery (3 repetitions in total). Flies were recorded, and videos were manually analyzed using VLC player (http://www.videolan.org/). Grooming activity (ethogram) was plotted as bins (1 bin, grooming event(s) per second). Ethograms were visualized using R (https://cran.r-project.org/). The ablation of 2^nd^ antennal segments did neither damage the head nor lead to fly mortality. Flies that died during the analysis window (7–15 dpa) were excluded.

### In-cell NAD-glo of dSarm proteins for NADase assay

#### Immunoprecipitation

S2 cells cell lysates (see above) were protein-quantified with the BCA protein assay and diluted to 500 ng/μl in ice-cold KHM buffer. Lysates were mixed with 20 μg/ml mouse anti-FLAG® M2 monoclonal antibody (Sigma-Aldrich F3165) and 50 μl/ml of pre-washed Pierce magnetic protein A/G beads (Thermo Fisher Scientific) and incubated overnight at 4 °C with rotation. After incubation, beads were washed 3x with KHM buffer and 1x with PBS and resuspended in 1 mg/ml BSA in PBS (with protease inhibitors).

#### NADase assays

A series of test assays were first performed to define appropriate test conditions. Optimized reaction conditions were as follows: 25 μl reactions (overall 1x PBS) contained 40 fmol/μl dSarm(E) protein together with 25 μM NAD^+^ ± 50 μM NMN. Reactions were kept on ice while being set up. Reactions were performed with the recombinant dSarm(E) still attached to beads and bead suspensions were thoroughly mixed prior to addition to the reactions. Constitutive (basal) NAD^+^ consumption was measured from reactions containing NAD^+^ alone as the difference between starting levels (0 mins) and levels remaining after incubating for between 80 and 120 min at 25 °C, and NAD^+^ consumption in the presence of 50 μM NMN was calculated after incubating for between 40 and 120 min (times were dependent on variant activity in each sample). Reactions were mixed once during the incubation to resuspend the beads. NAD^+^ levels were measured using the NAD/NADH-Glo™ assay. 5 μl aliquots of reaction were removed immediately after setting up (whilst still on ice), to obtain precise starting levels (0 min) in individual reactions, and again after the defined times listed above. Aliquots were then mixed with 2.5 μl of 0.4 M HCl, to stop the reaction, and neutralised by mixing with 2.5 μl 0.5 M Tris base after 10 min. Neutralised samples were subsequently diluted 1 in 50 in a buffer consisting of 50 % PBS, 25 % 0.4 M HCL, 25 % 0.5 M Tris base to bring the NAD^+^. NAD^+^ concentrations down to the linear range of detection for the NAD/NADH-Glo™ assay. 10 μl of the diluted sample was then mixed with 10 μl of NAD/NADH-Glo™ detection reagent on ice in wells of a 384-well white polystyrene microplate (Corning). Once all reactions had been set up the plate was moved to a GloMax® Explorer plate reader (Promega) and incubated for 40 min at 25 °C before reading for luminescence. NAD^+^ concentrations were determined from a standard curve generated from a dilution series of NAD^+^ and NAD^+^ consumption rates were converted to mol of NAD^+^ consumed per min per mol of dSarm(E) protein (mol/min/mol dSarm) ^53^. Individual data points for each separate protein preparation are the means of two or three technical replicates. No non-specific activity was detected on bead/antibody complexes in control immunoprecipitations using extracts from *Actin–Gal4* transfected S2 cells (based on *n* = 5).

### Replication

For all experiments, at least 3 biological replications were performed for each genotype and/or condition.

### Software and statistics

Image-J and photoshop was used to process wing and ORN pictures. Software for optogenetics is included in the optogenetics section. Graphpad prism 9 was used to perform all the statistical analysis. For tests applying a false discovery rate (FDR) correction, the adjusted p value we report is the q value.

## References

1. Riccomagno, M. M. & Kolodkin, A. L. Sculpting Neural Circuits by Axon and Dendrite Pruning, http://dx.doi.org/10.1146/annurev-cellbio-100913-013038 31, 779–805 (2015).

2. Neukomm, L. J. & Freeman, M. R. Diverse cellular and molecular modes of axon degeneration. Trends in Cell Biology 24, 515–523 (2014).

3. Coleman, M. P. & Höke, A. Programmed axon degeneration: from mouse to mechanism to medicine. Nature Reviews Neuroscience 2020 21:4 21, 183–196 (2020).

4. Mariano, V., Domínguez-Iturza, N., Neukomm, L. J. & Bagni, C. Maintenance mechanisms of circuit-integrated axons. Current Opinion in Neurobiology 53, 162–173 (2018).

5. Merlini, E., Coleman, M. P. & Loreto, A. Mitochondrial dysfunction as a trigger of programmed axon death. Trends in Neurosciences 45, 53–63 (2022).

6. Llobet Rosell, A. & Neukomm, L. J. Axon death signalling in Wallerian degeneration among species and in disease. Open Biology 9, 190118 (2019).

7. Waller, A. Experiments on the Section of the Glossopharyngeal and Hypoglossal Nerves of the Frog, and Observations of the Alterations Produced Thereby in the Structure of Their Primitive. Source: Philosophical Transactions of the Royal Society of London vol. 140 https://www.jstor.org/stable/pdf/108444.pdf?refreqid=excelsior%3A43333c1fb491b96c4dd2967aeff26769 (1850).

8. Gerdts, J., Summers, D. W., Sasaki, Y., DiAntonio, A. & Milbrandt, J. Sarm1-Mediated Axon Degeneration Requires Both SAM and TIR Interactions. Journal of Neuroscience 33, 13569–13580 (2013).

9. Osterloh, J. M. et al. dSarm/Sarm1 is required for activation of an injury-induced axon death pathway. Science (1979) 337, 481–484 (2012).

10. Essuman, K. et al. The SARM1 Toll/Interleukin-1 Receptor Domain Possesses Intrinsic NAD+Cleavage Activity that Promotes Pathological Axonal Degeneration. Neuron 93, 1334–1343.e5 (2017).

11. Figley, M. D. et al. SARM1 is a metabolic sensor activated by an increased NMN/NAD+ ratio to trigger axon degeneration. Neuron 109, 1118–1136.e11 (2021).

12. Gerdts, J., Brace, E. J., Sasaki, Y., Diantonio, A. & Milbrandt, J. Supplementary Materials for SARM1 activation triggers axon degeneration locally via NAD + destruction. Science (1979) 348, 453–458 (2015).

13. Hopkins, E. L., Gu, W., Kobe, B. & Coleman, M. P. A Novel NAD Signaling Mechanism in Axon Degeneration and its Relationship to Innate Immunity. Frontiers in Molecular Biosciences 8, 662 (2021).

14. Figley, M. D. & DiAntonio, A. The SARM1 axon degeneration pathway: control of the NAD + metabolome regulates axon survival in health and disease. Curr Opin Neurobiol 63, 59–66 (2020).

15. Gilley, J. & Coleman, M. P. Endogenous Nmnat2 Is an Essential Survival Factor for Maintenance of Healthy Axons. PLoS Biology 8, (2010).

16. Babetto, E., Beirowski, B., Russler, E., Milbrandt, J. & DiAntonio, A. The Phr1 Ubiquitin Ligase Promotes Injury-Induced Axon Self-Destruction. Cell Reports 3, 1422–1429 (2013).

17. Walker, L. J. et al. MAPK signaling promotes axonal degeneration by speeding the turnover of the axonal maintenance factor NMNAT2. Elife 6, (2017).

18. Loreto, A., di Stefano, M., Gering, M. & Conforti, L. Wallerian Degeneration Is Executed by an NMN-SARM1-Dependent Late Ca2+ Influx but Only Modestly Influenced by Mitochondria. Cell Reports 13, 2539–2552 (2015).

19. di Stefano, M. et al. A rise in NAD precursor nicotinamide mononucleotide (NMN) after injury promotes axon degeneration. Cell Death and Differentiation 22, 731–742 (2015).

20. di Stefano, M. et al. NMN Deamidase Delays Wallerian Degeneration and Rescues Axonal Defects Caused by NMNAT2 Deficiency In Vivo. Curr Biol 27, 784–794 (2017).

21. Loreto, A. et al. Neurotoxin-mediated potent activation of the axon degeneration regulator SARM1. Elife 10, (2021).

22. Bratkowski, M. et al. Structural and Mechanistic Regulation of the Pro-degenerative NAD Hydrolase SARM1. (2020) doi:10.1016/j.celrep.2020.107999.

23. Zhao, Z. Y. et al. A Cell-Permeant Mimetic of NMN Activates SARM1 to Produce Cyclic ADP-Ribose and Induce Non-apoptotic Cell Death. iScience 15, 452 (2019).

24. Jiang, Y. et al. The NAD+-mediated self-inhibition mechanism of pro-neurodegenerative SARM1. Nature 2020 588:7839 588, 658–663 (2020).

25. Angeletti, C. et al. SARM1 is a multi-functional NAD(P)ase with prominent base exchange activity, all regulated by multiple physiologically-relevant NAD metabolites. iScience 103812 (2022) doi:10.1016/J.ISCI.2022.103812.

26. Galeazzi, L. et al. Identification of nicotinamide mononucleotide deamidase of the bacterial pyridine nucleotide cycle reveals a novel broadly conserved amidohydrolase family. Journal of Biological Chemistry 286, 40365–40375 (2011).

27. Sasaki, Y., Nakagawa, T., Mao, X., DiAntonio, A. & Milbrandt, J. NMNAT1 inhibits axon degeneration via blockade of SARM1-mediated NAD+ depletion. Elife 5, (2016).

28. Ruan, K., Zhu, Y., Li, C., Brazill, J. M. & Zhai, R. G. Alternative splicing of Drosophila Nmnat functions as a switch to enhance neuroprotection under stress. Nature Communications 6, 10057 (2015).

29. Xiong, X. et al. The Highwire Ubiquitin Ligase Promotes Axonal Degeneration by Tuning Levels of Nmnat Protein. PLoS Biology 10, 1–18 (2012).

30. Neukomm, L. J. et al. Axon Death Pathways Converge on Axundead to Promote Functional and Structural Axon Disassembly. Neuron 95, 78–91.e5 (2017).

31. Gossmann, T. I. et al. NAD+ biosynthesis and salvage – a phylogenetic perspective. The FEBS Journal 279, 3355–3363 (2012).

32. Gerdts, J., Summers, D. W., Milbrandt, J. & DiAntonio, A. Axon Self-Destruction: New Links among SARM1, MAPKs, and NAD+ Metabolism. Neuron 89, 449–460 (2016).

33. Hsu, J. M. et al. Injury-Induced Inhibition of Bystander Neurons Requires dSarm and Signaling from Glia. Neuron 109, 473–487.e5 (2021).

34. Fang, Y., Soares, L., Teng, X., Geary, M. & Bonini, N. M. A novel drosophila model of nerve injury reveals an essential role of Nmnat in maintaining axonal integrity. Current Biology 22, 590–595 (2012).

35. Neukomm, L. J., Burdett, T. C., Gonzalez, M. A., Züchner, S. & Freeman, M. R. Rapid in vivo forward genetic approach for identifying axon death genes *in Drosophila*. Proceedings of the National Academy of Sciences 111, 9965–9970 (2014).

36. Paglione, M., Llobet Rosell, A., Chatton, J. Y. & Neukomm, L. J. Morphological and Functional Evaluation of Axons and their Synapses during Axon Death in Drosophila melanogaster. J. Vis. Exp 60865 (2020) doi:10.3791/60865.

37. Rücker, E. et al. Rapid evaluation and optimization of recombinant protein production using GFP tagging. Protein Expr Purif 21, 220–223 (2001).

38. Amici, A. et al. Synthesis and Degradation of Adenosine 5Ø-Tetraphosphate by Nicotinamide and Nicotinate Phosphoribosyltransferases. Cell Chemical Biology 24, 553–564.e4 (2017).

39. Zamporlini, F. et al. Novel assay for simultaneous measurement of pyridine mononucleotides synthesizing activities allows dissection of the NAD(+) biosynthetic machinery in mammalian cells. FEBS J 281, 5104–5119 (2014).

40. van der Velpen, V. et al. Sex-specific alterations in NAD+ metabolism in 3xTg Alzheimer’s disease mouse brain assessed by quantitative targeted LC-MS. Journal of Neurochemistry 159, 378–388 (2021).

41. Hampel, S., Franconville, R., Simpson, J. H. & Seeds, A. M. A neural command circuit for grooming movement control. Elife 4, (2015).

42. Seeds, A. M. et al. A suppression hierarchy among competing motor programs drives sequential grooming in Drosophila. Elife 3, e02951 (2014).

43. Gu, W. et al. Crystal structure determination of the armadillo repeat domain of Drosophila SARM1 using MIRAS phasing. urn:issn:2053-230 77, 364–373 (2021).

44. Geisler, S. et al. Gene therapy targeting SARM1 blocks pathological axon degeneration in mice. J Exp Med 216, 294–303 (2019).

45. Zhai, R. G. et al. Drosophila NMNAT maintains neural integrity independent of its NAD synthesis activity. PLoS Biology 4, 2336–2348 (2006).

46. Port, F. & Bullock, S. L. Augmenting CRISPR applications in Drosophila with tRNA-flanked sgRNAs. Nature Methods 13, 852–854 (2016).

47. Sasaki, Y. et al. Nicotinic acid mononucleotide is an allosteric SARM1 inhibitor promoting axonal protection. Experimental Neurology 345, 113842 (2021).

48. Lehmann, S., Loh, S. H. Y. & Martins, L. M. Enhancing NAD + salvage metabolism is neuroprotective in a PINK1 model of Parkinson’s disease. Biol Open 6, 141–147 (2017).

49. Yoshino, J., Baur, J. A. & Imai, S. ichiro. NAD+ Intermediates: The Biology and Therapeutic Potential of NMN and NR. Cell Metabolism 27, 513–528 (2018).

50. Liu, L. et al. Quantitative Analysis of NAD Synthesis-Breakdown Fluxes. Cell Metabolism 27, 1067–1080.e5 (2018).

51. Izadifar, A. et al. Axon morphogenesis and maintenance require an evolutionary conserved safeguard function of Wnk kinases antagonizing Sarm and Axed. Neuron 109, 2864–2883.e8 (2021).

52. Ji, H., Sapar, M. L., Sarkar, A., Wang, B. & Han, C. Phagocytosis and self-destruction break down dendrites of Drosophila sensory neurons at distinct steps of Wallerian degeneration. Proceedings of the National Academy of Sciences 119, e2111818119 (2022).

53. Gilley, J. et al. Enrichment of SARM1 alleles encoding variants with constitutively hyperactive NADase in patients with ALS and other motor nerve disorders. Elife 10, (2021).

