## Supplementary figures and images for "The NAD^+^ precursor NMN activates dSarm to trigger axon degeneration in *Drosophila*"

### Supplementary Figure S1

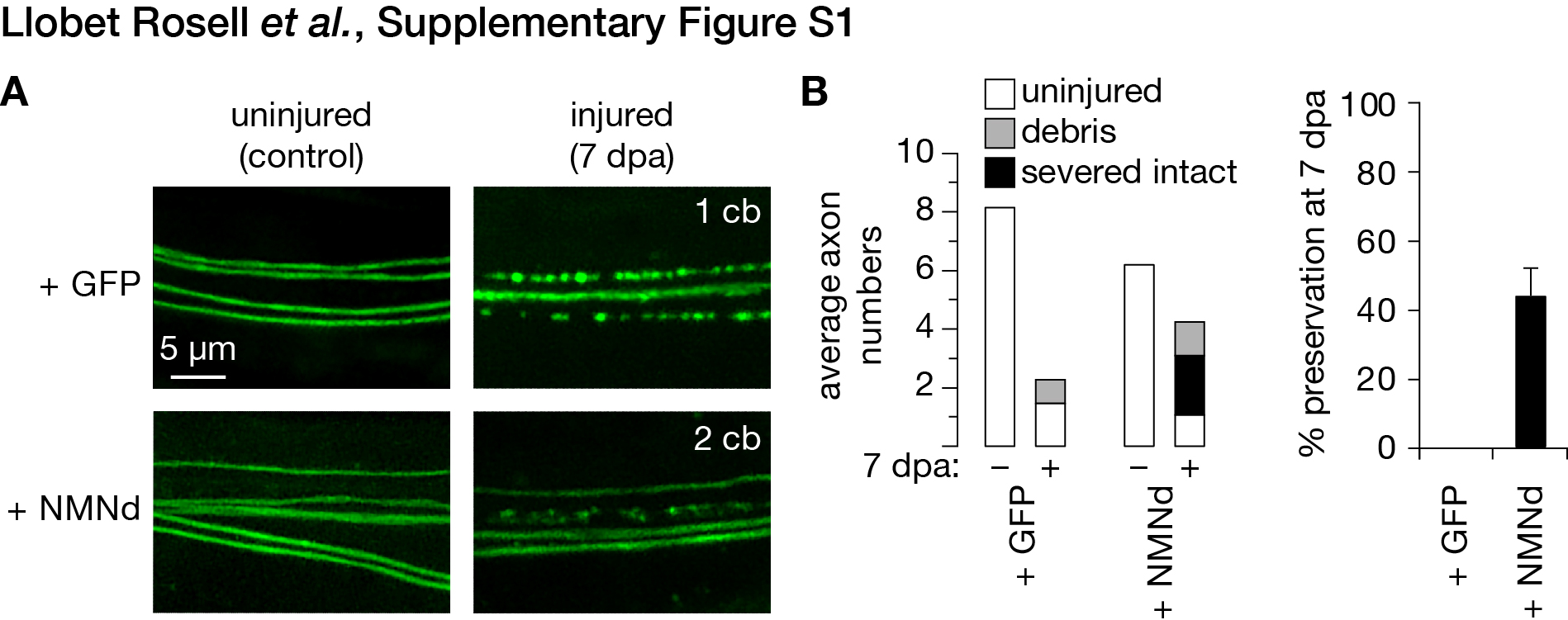

### Supplementary Figure S2

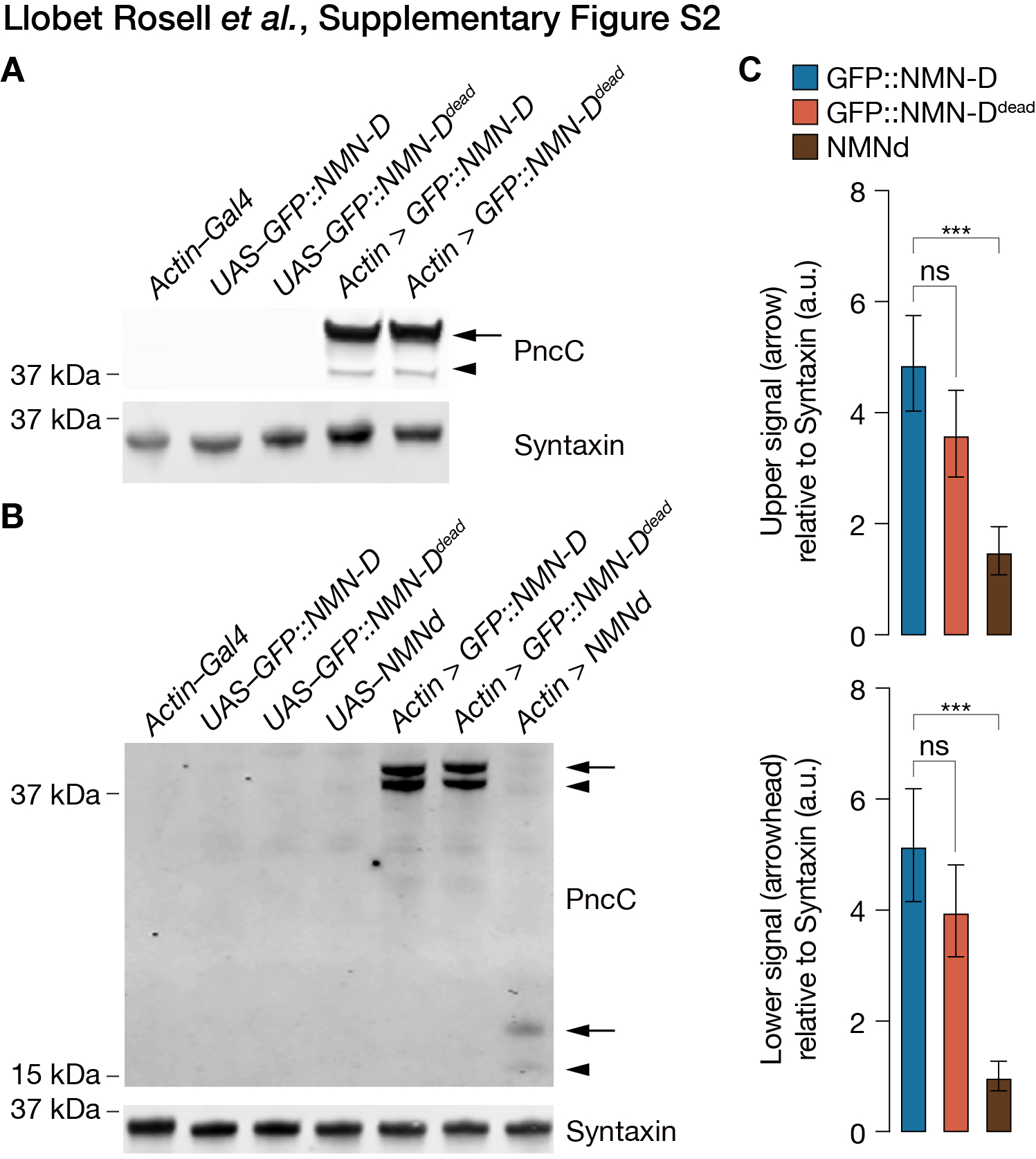

### Supplementary Figure S3

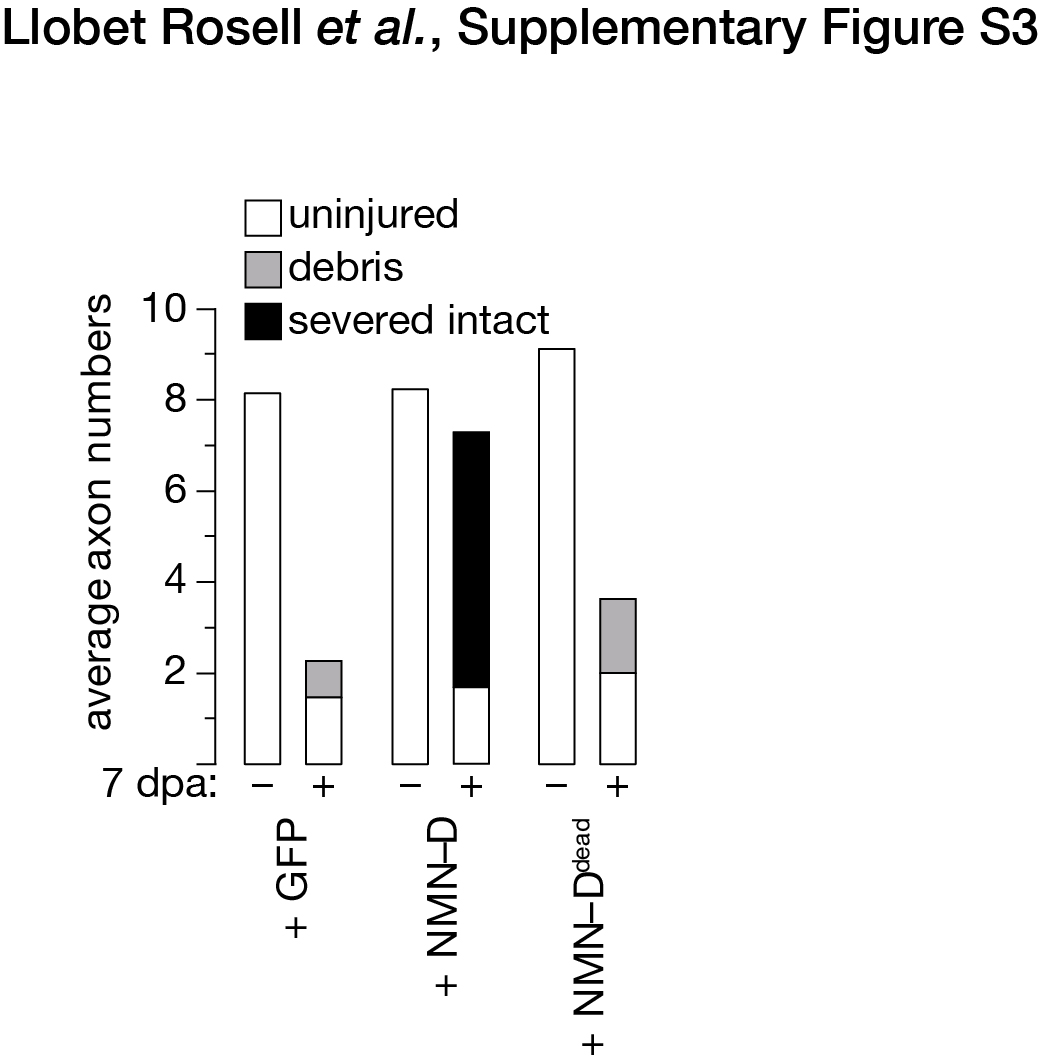

### Supplementary Figure S4

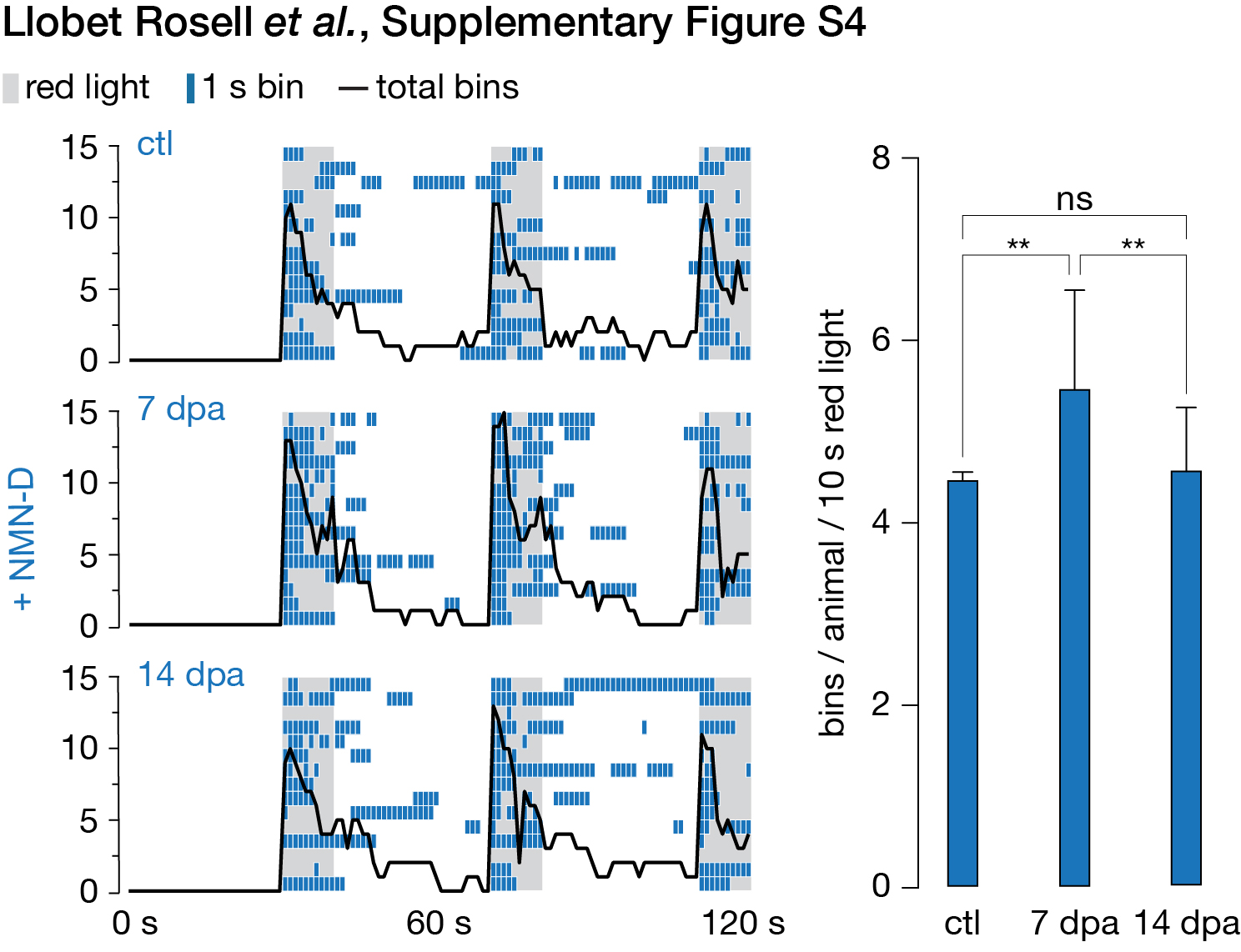

### Supplementary Figure S5

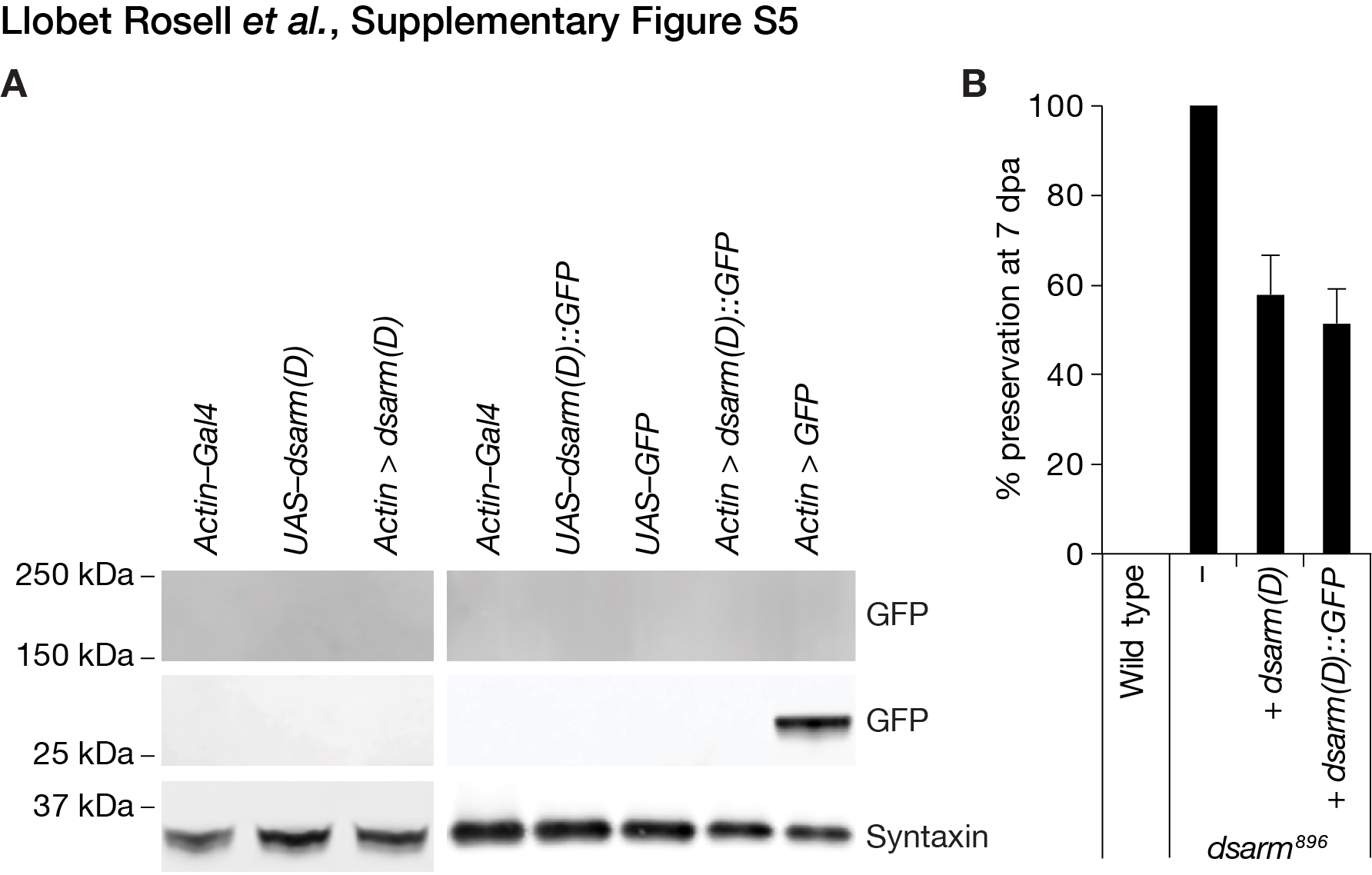

### Supplementary Figure S6

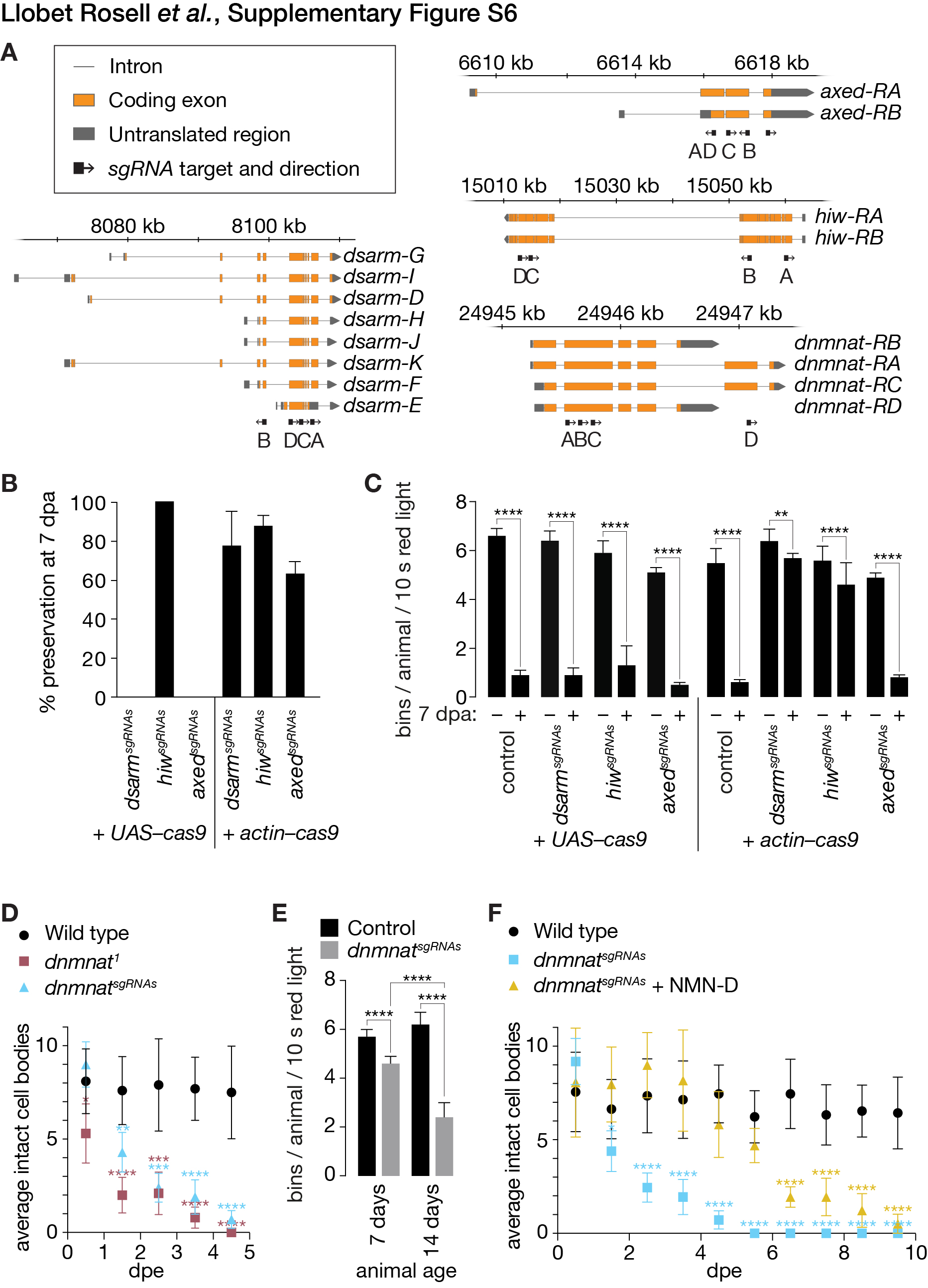

### Supplementary Table S2

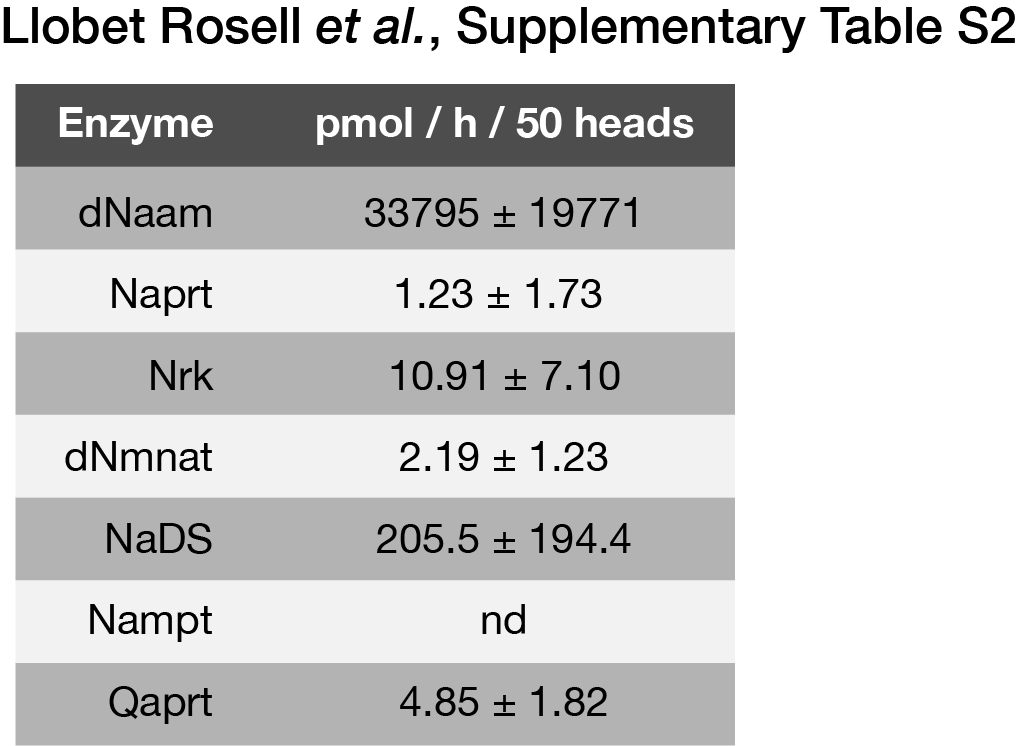
